# Super-resolution imaging reveals the evolution of higher-order chromatin folding in early carcinogenesis

**DOI:** 10.1101/672105

**Authors:** Jianquan Xu, Hongqiang Ma, Hongbin Ma, Wei Jiang, Meihan Duan, Shimei Zhao, Chenxi Gao, Eun-Ryeong Hahm, Santana M. Lardo, Kris Troy, Ming Sun, Reet Pai, Donna B Stolz, Shivendra Singh, Randall E Brand, Douglas J. Hartman, Jing Hu, Sarah J. Hainer, Yang Liu

## Abstract

Aberrant chromatin structure is a hallmark in cancer cells and has long been used for clinical diagnosis of cancer. However, underlying higher-order chromatin folding during malignant transformation remains elusive, due to the lack of molecular scale resolution. Using optimized stochastic optical reconstruction microscopy (STORM) for pathological tissue (PathSTORM), we uncovered a gradual decompaction and fragmented higher-order chromatin folding throughout all stages of carcinogenesis in multiple tumor types, even prior to the tumor formation. Our integrated imaging, genomic, and transcriptomic analyses reveal the functional consequences in enhanced formation of transcription factories, spatial juxtaposition with relaxed nanosized chromatin domains and impaired genomic stability. We also demonstrate the potential of imaging higher-order chromatin decompaction to detect high-risk precursors that cannot be distinguished by conventional pathology. Taken together, our findings reveal the gradual decompaction and fragmentation of higher-order chromatin structure as an enabling characteristic in early carcinogenesis to facilitate malignant transformation, which may improve cancer diagnosis, risk stratification, and prevention.

**SIGNIFICANCE:** Genomic DNA is folded into a higher-order structure that regulates transcription and maintains genomic stability. Although much progress has been made on understanding biochemical characteristics of epigenetic modifications in cancer, the higher-order folding of chromatin structure remains largely unknown. Using optimized super-resolution microscopy, we uncover de-compacted and fragmented chromatin folding in tumor initiation and stepwise progression in multiple tumor types, even prior to the presence of tumor cells. This study underlines the significance of unfolding higher-order chromatin structure as an enabling characteristic to promote tumorigenesis, which may facilitate the development and evaluation of new preventive strategies. The potential of imaging higher-order chromatin folding to improve cancer detection and risk stratification is demonstrated by detecting high-risk precursors that cannot be distinguished by conventional pathology.

## INTRODUCTION

Aberrant chromatin structure visualized under conventional light microscopy is one of the most universal and striking characteristics in cancer cells (Brock et al., 2007) and has long been used to diagnose cancer (Zink et al., 2004). Despite its clinical significance, the molecular underpinning of aberrant chromatin structure in cancer cells remains elusive (Fischer et al., 2010; Reddy and Feinberg, 2013). The fundamental unit of chromatin is the nucleosome which is further packaged into higher-order structure, organized into condensed, transcriptionally repressed heterochromatin and open, transcriptionally active euchromatin domains. In particular, constitutive heterochromatin—the most condensed form of chromatin, enriched for trimethylated histone H3 lysine 9 (H3K9me3) and repetitive sequence—protects genomic integrity and stability (Janssen et al., 2018).

Many studies showed that defective heterochromatin results in genomic instability, an “enabling characteristic” for cells to acquire hallmarks of cancer (Hanahan and Weinberg, 2011). Double knockout mice of H3K9 methyltransferases Suv39h1 and Suv39h2 exhibit chromosome instability and increased risk of tumor formation (Peters et al., 2001). Large organized heterochromatin H3K9 modifications were substantially lost in several cancer cell lines (Wen et al., 2009) and in cells undergoing epithelial-to-mesenchymal transition (McDonald et al., 2011), a crucial mechanism in cancer development. Loss of heterochromatin reader protein HP1 causes chromosome segregation and is associated with cancer progression (Dialynas et al., 2008). Depletion of a chromatin modifier, G9a, results in the development of tumors in mice that are more aggressive tumors and genomically unstable (Avgustinova et al., 2018). At the genomic level, satellite repeats (enriched with H3K9me3) are required for heterochromatin formation and accurate chromosome segregation (Saksouk et al., 2015).

Although the causal relationship between dysregulated heterochromatin function and increased genomic instability is a well-recognized mechanism to promote cancer progression, it is largely inferred by biochemical analysis of chromatin-associated proteins and DNA sequences. The underlying molecular-scale chromatin structures for dysregulated heterochromatin function in tumorigenesis from tumor initiation to stepwise progression remains largely unknown. This is, in part, due to the lack of sufficient resolution to visualize molecular scale higher-order chromatin structure, which is below the resolution limit of a conventional light microscope. Several important questions remain unanswered. What is the characteristic molecular-scale chromatin abnormality in cancer cells? Coarse aggregation of heterochromatin has been a well-documented microscopic feature diagnostic of many cancers (Zink et al., 2004), but numerous biological studies reported the loss of heterochromatin-associated proteins and post-translational marks in cancer cells which cannot explain the aggregated heterochromatin structure (Avgustinova et al., 2018; Janssen et al., 2018; Peters et al., 2001). Aberrant chromatin structure is a characteristic of cancer cells when normal cells have already turned into tumor, but what happens in early carcinogenesis when cells still appear normal prior to tumor formation? Is the structural disruption a one-hit event or a gradual evolving process throughout carcinogenesis? Is there a common feature in chromatin structure underlying all stages of carcinogenesis, independent of cancer-driven molecular pathways and tumor types? Answering these questions will have significant clinical implications to improve early detection and/or cancer risk stratification. For example, in precursors that are indistinguishable by pathologists, whether those high-risk precursors undergoing aggressive progression to cancer exhibit distinct heterochromatin structure from those with low risk?

Recent advances in super-resolution fluorescence microscopy has revolutionized biological imaging by overcoming the diffraction barrier that limits the resolution of conventional light microscope to be ∼250 nm (Sigal et al., 2018). In particular, (direct) stochastic optical reconstruction microscopy [(d)STORM] (Enderlein, 2015; Rust et al., 2006) has shown great promise in advancing our fundamental understanding of *in situ* nanoscale chromatin structure down to the level of nucleosome clusters (or nanodomains) at the scale of ∼30 nm in cultured cells and small organisms (e.g., *Drosophila*) (Beliveau et al., 2015; Boettiger et al., 2016; Prakash et al., 2015; Ricci et al., 2015; Xu et al., 2018). However, characteristic higher-order chromatin structure throughout tumorigenesis has not been fully characterized on pathological tissue with well-preserved spatial context of tissue architecture, in part due to challenges in imaging pathological tissue such as autofluorescence, stronger scattering, and non-uniform background.

We optimized a STORM-based super-resolution imaging method for robust and high-speed super-resolution imaging of higher-order chromatin structure on pathological tissues. Super-resolution imaging revealed significant fragmentation in DNA folding and heterochromatin decompaction at the earliest stage of carcinogenesis even in normal-appearing tissue at risk for tumorigenesis. Integrated genomic and transcriptomic analysis also suggested an opening of chromatin structure and revealed that disrupted heterochromatin occurs mostly in the satellite repeat regions of the genome with increased gene expression. We also observed enhanced formation of transcription factories juxtaposed with relaxed chromatin domains and impaired genomic stability as functional consequences after we disrupt heterochromatin structure. Furthermore, we found a gradual decompaction of heterochromatin structure throughout all stages of tumorigenesis, which is also a common feature independent of molecular pathways from multiple tumor types. Finally, we showed the potential of super-resolution imaging of heterochromatin decompaction to detect high-risk precursors that cannot be distinguished by conventional pathology. Together, our finding of heterochromatin decompaction at the nucleosome level underlies its importance in early carcinogenesis and opens a new avenue for improving cancer diagnosis, risk stratification, and facilitating the development and evaluation of new preventive strategies.

## RESULTS

### PathSTORM visualizes higher-order chromatin structure in pathological tissue

The super-resolved imaging capability of STORM is largely based on precise localization of sparsely distributed single fluorescent emitters at nanometer precision, and generally achieves best performance in thin cultured cells. But stronger scattering and autofluorescence in tissue often produce high and non-uniform background that can significantly degrade image resolution and introduce artifacts (Deschout et al., 2014). We optimized STORM for imaging pathological tissue (referred to as PathSTORM), especially for those formalin-fixed paraffin-embedded (FFPE) pathological tissue – the most common form of preserved pathological specimens. We obtained high-fidelity super-resolution images of higher-order chromatin structure on FFPE tissue section with the following three approaches: (1) optical clearing and index matching to reduce background, (2) a new extreme value-based emitter recovery to correct residual heterogeneous background, and (3) a computationally effective localization method to improve localization precision of overlapping emitters (Ma et al., 2019), which are shown in Supplementary Fig S1.

We then validated the ability of PathSTORM to visualize higher-order chromatin structures at different epigenomic states *in vivo* in normal intestinal epithelial tissue (Supplementary Fig. S2). The super-resolution images of heterochromatin and euchromatin show distinct and heterogeneous groups of nucleosome clusters at the scale of tens of nanometers, serving as the building blocks for higher-order chromatin structure. In particular, heterochromatin forms highly condensed large nanoclusters, while euchromatin exhibits more uniform or spatially diffuse nanoclusters, as indicated by the green arrows in Fig. S2. These results are consistent with previously reported higher-order chromatin structure on the *in vitro* mammalian cell cultures (Ricci et al., 2015; Xu et al., 2018). We further confirmed the observed higher-order chromatin structure by comparing the reconstructed super-resolution images with those from the ultra-thin frozen tissue section. Ultra-thin frozen tissue section (700 nm-thick) has been used as a strategy to achieve low and uniform background with standard STORM imaging condition (Sigal et al., 2015). As shown in Supplementary Fig. S3, the chromatin structure from FFPE tissue shows similar structural features with similar cluster size and density. This result quantitatively validated the ability of PathSTORM to achieve similar resolution of higher-order chromatin structures on pathological tissue compared to the standard STORM.

### PathSTORM reveals higher-order heterochromatin decompaction and fragmented DNA folding in early carcinogenesis in a mouse model

Next, to visualize the changes of higher-order heterochromatin structure in carcinogenesis, we used a well-established mouse model of intestinal tumorigenesis – *Apc*^Min/+^ mouse where a mutation in the tumor suppressor gene *adenomatous polyposis coli* (*Apc*) causes spontaneous development of multiple intestinal neoplasia (Min) and closely mimics familial adenomatous polyposis in human colorectal neoplasia (Su et al., 1992). Heterochromatin was fluorescently labeled with H3K9me3 that largely overlaps with the most condensed regions of DNA (stained with DAPI) in the epithelial tissue (see Supplementary Fig. S4). Figure 1 shows the hematoxylin and eosin (H&E)-stained histology images (A1-C1), conventional wide-field fluorescence (A2-C2, A3-C3), and corresponding super-resolution images of heterochromatin in three types of tissue – (a) normal intestinal epithelial cell nuclei of wild-type mice, (b) normal-appearing epithelial cell nuclei of intestinal mucosa at risk for tumorigenesis from 6-week *Apc*^Min/+^ mice without any visible tumor and (c) tumor cell nuclei (adenomatous polyps) from 12-week *Apc*^Min/+^ mice. The conventional wide-field fluorescence images (Figs. 1(A2-C2, A3-C3)) show large and dense heterochromatin foci in all three types of tissue. In contrast, super-resolution images reveal that each of the large heterochromatin foci is formed by groups of nucleosome nanoclusters. In normal cell nuclei from wild-type mice, their nanoclusters are highly compact; in 6-week *Apc*^Min/+^ mice at the early stage of carcinogenesis when tissue still appears normal, surprisingly, the nucleosome-level nanoclusters already become smaller and spatially segregated (Figs. 1(B4-B6)), which are even smaller in the tumor cells (adenoma) in 12-week *Apc*^Min/+^ mice (Figs. 1(C4-C6)). This result clearly shows fragmentation of heterochromatin foci and nucleosome-level decompaction of higher-order heterochromatin structure. Such disruption in higher-order heterochromatin structure cannot be easily observed in conventional fluorescence images.

**Figure 1.**
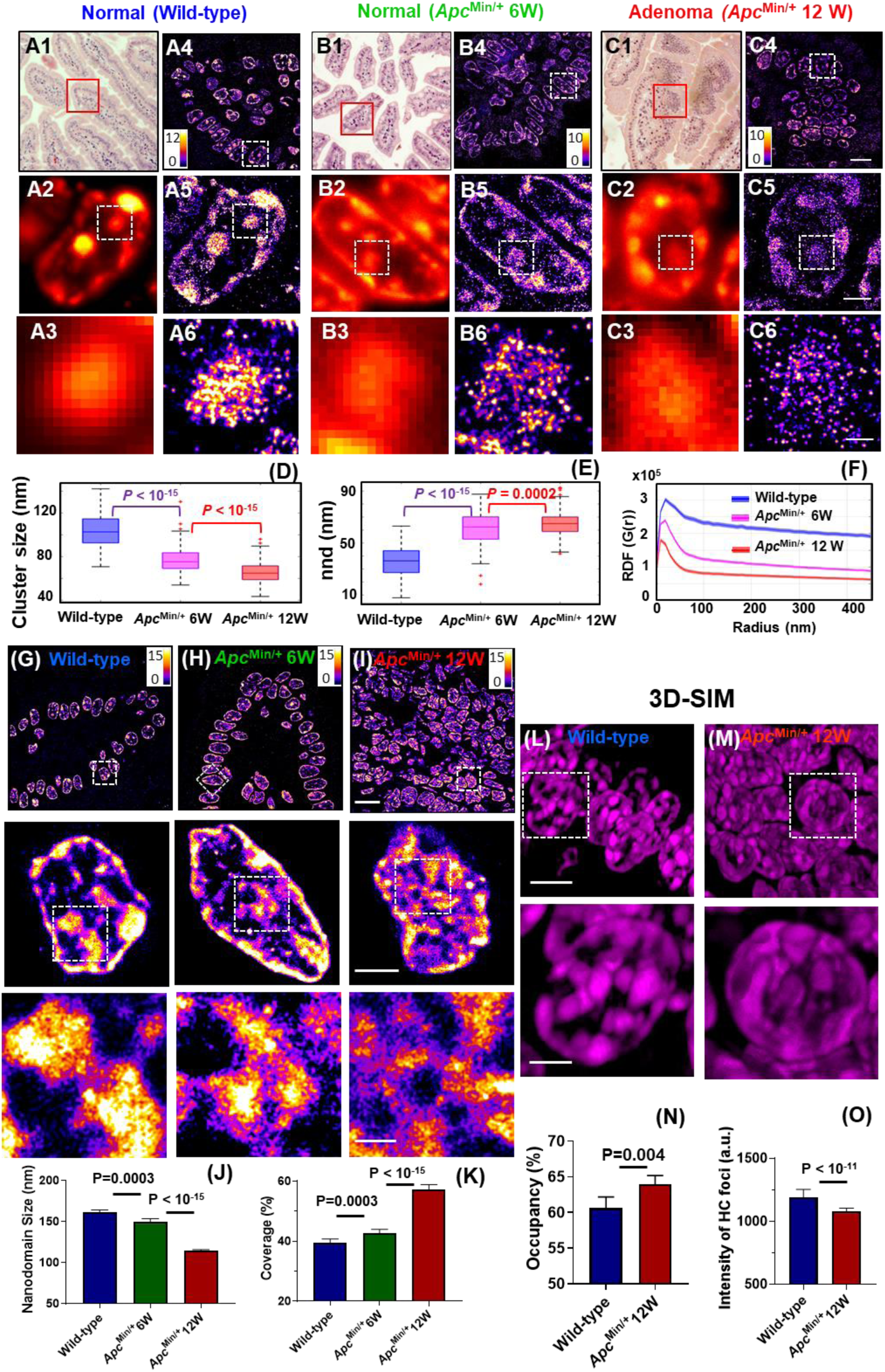
Super-resolution imaging of H3K9me3 and DNA shows the decompaction of heterochromatin and fragmented DNA folding in carcinogenesis in *Apc*^Min/+^ mouse model. (**A1-C1**) The H&E-stained pathology images and (**A4-C4**) STORM-based super-resolution images of H3K9me3-dependent heterochromatin from the red boxes in (A1-C1) from normal tissue from wild-type, histologically normal-appearing tissue from 6-week *Apc*^Min/+^ and tumor (adenoma) from 12-week *Apc*^Min/+^ mice. Scale bar: 10µm. (**A2-C2**) Conventional wide-field fluorescence images and (**A5-C5**) corresponding STORM images of heterochromatin from a single nucleus. (**A3-C3**) and (**A6-C6**) are progressively zoomed regions of (A2-C2) and (A5-C5). Scale bar in (A5-C5): 2µm; scale bar in (A6-C6): 500nm. Scale bars, 10 µm, 2 µm and 500 nm in the original and magnified images, respectively (**D-E**) The box-and-whisker plots of the H3K9me3 cluster size (C) and nearest neighbor distance (nnd) between clusters (outliers: “+”). Over 150 cell nuclei were analyzed for each group. *P* values were determined using Mann-Whitney test. (**F**) Average radial distribution function (RDF) for all nuclei in each group. The solid curve shows the average RDF from all measured nuclei and the shaded area shows the standard error. (**G-I**) The STORM images of DNA folding from normal cells from wild-type, histologically normal-appearing cells from 6-week *Apc*^Min/+^ and tumor cells from 12-week *Apc*^Min/+^ mice. Scale bars: 10 µm, 2 µm and 500 nm in the original and magnified images, respectively. (**J-K**) The statistical average of DNA nanodomain size and the percentage of occupied DNA domains within each nucleus in normal cells from wild-type mice, from 6-week *Apc*^Min/+^ mice and tumor cells from 12-week *Apc*^Min/+^ mice. (**L-M**) The 3D-SIM images of DAPI-stained DNA folding in normal cells from wild-type mice and tumor cells from 12-week *Apc*^Min/+^ mice. Scale bars: 5 µm, and 2 µm in the original and magnified images, respectively. (**N**) The occupany of DNA defined by the total volume of DNA over the entire 3D volume of each nucleus, calcuated from the 3D-SIM images. (**O**) The average fluorescence intensity from each of the condensed region of heterochromatin (HC) foci in 3D-SIM images.

We then quantified the structural disruption of heterochromatin using two different methods – Gaussian mixed model clustering (Ma et al., 2017) and radial distribution function (RDF) (a.k.a. pair-correlation function) (Caetano et al., 2015; Xu et al., 2018). The former quantifies the size of nucleosome clusters (or the building blocks that form the large heterochromatin foci); the latter provides a global overview for all heterochromatin structures existing on multiple length scales. Figures 1(D-E) show that, during carcinogenesis, there is a progressively decreased size of heterochromatin nanoclusters and increased nearest neighbor distance (nnd) with a high statistical significance (*P*<10^−15^) (>150 cell nuclei analyzed). The RDF distribution in Fig. 1F also shows a progressively narrower distribution and smaller correlation length, consistent with our observed progressive disruption of heterochromatin foci in Figs. 1(A-C). Moreover, we did not observe a significant difference in the normal epithelial cell nuclei between 6-week and 12-week wild-type mice (see Supplementary Fig. S5), suggesting that our observed higher-order structural disruption in heterochromatin is not due to a mere age difference.

To further evaluate the disruption of heterochromatin structure, we also performed super-resolution imaging on DNA labeled with a fluorophore that directly binds to nucleic acids. As shown in Figs. 1(G-I), the STORM images of DNA also reveal a more fragmented and relaxed DNA folding from normal-appearing cells at risk for tumorigenesis (6-week *Apc*^Min/+^ mice) compared to normal cells from wild-type mice. In tumor cells from 12-week *Apc*^Min/+^ mice, the relaxed and fragmented DNA folding becomes much more prominent. Quantitative analysis of STORM images shows a progressively reduced size of DNA nanoclusters in 6-week and 12-week *Apc*^Min/+^ mice (Fig. 1J). The area of the nucleus covered by DNA is significantly increased along with tumorigenesis (*P* = 0.0003 and *P* < 10^−15^ for normal cells at risk for tumorigenesis and tumor cells, respectively) (Fig. 1K). These results are consistent with the above-observed decompaction of H3K9me3-labeled heterochromatin. The fragmented DNA feature and decreased nanodomain size indicate the relaxation of chromatin folding in the nucleus. In addition, we confirmed that our observed difference in higher-order chromatin structure between normal and tumor cells is not due to the difference in cell cycle (Supplementary Fig. S6).

We further examined 3D heterochromatin structure (stained with DAPI) of normal cells from wild-type mice and tumor cells from 12-week *Apc*^Min/+^ mice using another super-resolution technique – 3D structured illumination microscopy (3D-SIM) (Figs. 1L-M). As SIM has a lower resolution (∼120 nm) compared to that of PathSTORM (∼30 nm), the fragmented DNA nanoclusters observed in PathSTORM cannot be easily seen in the 3D-SIM images. Instead, we observed more enlarged heterochromatin foci, as evidenced by increased DNA occupancy (defined as the percentage of volumes with DNA over the entire 3D volume of the cell nucleus) in tumor cells (Fig. 1N). Further, as shown in Fig. 1O, the fluorescence intensity from the dense heterochromatin foci of tumor cells is also significantly lower compared to that of normal cells from wild-type mice. The enlarged heterochromatin foci and reduced fluorescent intensity from the foci collectively suggest structural decompaction. This observation of enlarged heterochromatin foci in tumor cells is largely in line with the common cytologic diagnostic criteria of “coarse aggregates of heterochromatin” in cancer cells under bright-field microscope (Zink et al., 2004).

### Disrupted heterochromatin coincides with reduced occupancy of H3K9me3 at regions of satellite repeats

To identify the specific genomic regions affected by the disrupted heterochromatin structure, using CUT&RUN (Hainer et al., 2019; Skene and Henikoff, 2017), we profiled native (no crosslinking) chromatin structure of intestinal epithelial cells brushed from normal mouse intestine of wild-type and age-matched *Apc*^Min/+^ mice at 6 weeks, which is in early carcinogenesis when tissue still appears histologically normal. CUT&RUN is a newly developed genome-scale protein localization technique that serves as an alternative to chromatin immunoprecipitation (ChIP) but provides lower background signal. We profiled the occupancy of H3K9me3 and total histone H3, as well as a control lacking a primary antibody (but still including the proteinA-MNase to control for nonspecific fragment digestion and release) referred to as “no antibody”. As shown in Figs. 2(A-B), when we average the signal for H3K9me3 over previously described H3K9me3 genomic locations, we observed an enrichment of H3K9me3 relative to surrounding regions, while the “no antibody” control shows low background signal. In addition, the substantial reduction in H3K9me3 levels marking heterochromatin is accompanied by increased levels of H3K4me3 (an euchromatin marker) at promoter regions, suggesting more open chromatin (Supplementary Fig. S7).

**Figure 2.**
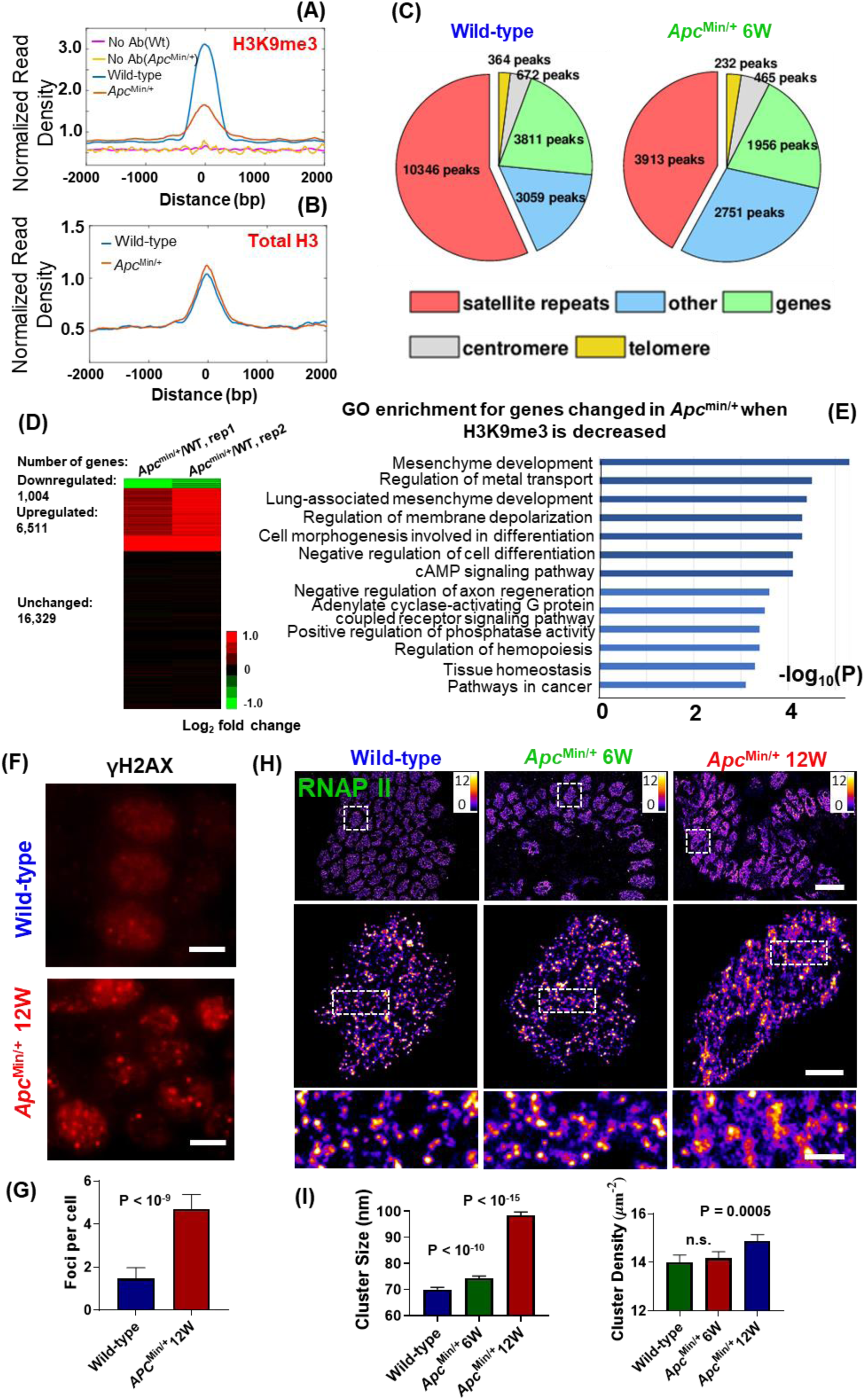
Disrupted heterochromatin structure leads to increased transcription and genomic instability. (**A-B**) The average enrichment of H3K9me3 and total H3 from previously described H3K9me3 enriched genomic regions, and (**C**) pie charts that show the genomic distribution of peaks that were enriched by H3K9me3, from normal-appearing intestinal tissue from wild-type mice and age-matched *Apc*^Min/+^ mice at 6 weeks. The genomic distribution in the pie chart consists of satellite repeats, genes, centromere, telomere and other genomic regions (including intergenic, other repeat regions that are unassigned to a type of repeat and non-annotated regions of the genome). The average from two duplicate experiments is shown in (A-B). (**D**) The heatmap of two duplicate experiments shows the differential gene expression of epithelial tissue between *Apc*^Min/+^ mice at 6 weeks and age-matched wild-type mice. Red color shows up-regulated genes, green color shows down-regulated genes and black color shows genes with no change in expression. (**E**) The gene ontology (GO) analysis for overlapping up-regulated genes with reduced occupancy of H3K9me3. (**F**) γH2AX immunofluorescence staining in wild-type mice and *Apc*^Min/+^ mice at 12 weeks. (**G**) The number of γH2AX foci for each group. (**H**) The STORM images of active RNAP II from normal cells from wild-type, histologically normal-appearing cells from 6-week *Apc*^Min/+^ and tumor cells from 12-week *Apc*^Min/+^ mice. Scale bars: 10 µm, 2 µm and 500 nm in the original and magnified images, respectively. (**I**) Statistical analysis of the active RNAPII cluster size and density for each group. Over 300 cell nuclei were analyzed for each group. *P-*values were determined using non-parametric Mann-Whitney test.

As shown in the pie chart in Fig. 2(C), the majority of H3K9me3 peaks identified in wild-type mice are within satellite repeats, as previously shown (Bulut-Karslioglu et al., 2014; Elsasser et al., 2015; Maze et al., 2011). In comparison, in 6-week *Apc*^Min/+^ mice, the occupancy of H3K9me3 is significantly reduced, with less peaks identified overall and the most significant reduction occurring at regions of satellite repeats (from 10346 to 3913 peaks), with the occupancy of total H3 protein remaining unchanged. This result not only supports our imaging finding of disrupted heterochromatin structure (reduced size of H3K9me3 nanoclusters) identified by PathSTORM; but also reveals that the disrupted heterochromatin mostly affects regions of satellite repeats in the genome of normal-appearing cells at the early-stage carcinogenesis.

To explore the functional consequence of the observed disrupted heterochromatin structure, we performed genome-wide total RNA-seq on the same sets of mouse tissue. As shown in Fig. 2D, differential gene expression between the 6-week *Apc*^Min/+^ mice and wild-type mice shows the majority of genes with altered expression are upregulated, which is in line with increased overall transcription and open chromatin structure during tumorigenesis (Brock et al., 2007; Timp and Feinberg, 2013). To identify the changes in gene expression directly altered by changes in H3K9me3 levels, we integrated CUT&RUN and RNA-seq analysis by assessing the gene expression changes of those genes that had reduced H3K9me3 in 6-week *Apc*^Min/+^ mice. Out of the 1855 genes that had reduced H3K9me3 levels, 820 genes showed a 2-fold or greater increase in gene expression (∼44% of genes with H3K9me3 reduction). As shown in Fig. 2E, Gene Ontology (GO) enrichment analysis identifies that the most significantly enriched shared pathway in this gene set is mesenchymal development (*P* < 10^−5^), a process leading to acquisition of invasiveness and malignancy in epithelial cancer cells (McDonald et al., 2011). GO enrichment analysis of the genes associated with increased H3K4me3 occupancy in 6-week *Apc*^Min/+^ mice identifies the most significantly enriched pathways as cellular response to DNA damage stimulus and mitotic cell cycle process (*P* < 10^−8^), suggesting impaired genomic stability as the potential functional consequence of open chromatin (Supplementary Fig. S7).

### Disrupted heterochromatin structure leads to genomic instability and enhanced formation of transcription factories

The above genomic and transcriptomic analysis suggests genomic instability and increased gene expression as a result of disrupted heterochromatin structure in early carcinogenesis. As shown in Figs. 2(F-G), we observed a significantly increased number of γ-H2AX foci (*P* < 10^−9^)—a marker for DNA double-strand breaks—in *Apc*^Min/+^ mice, suggesting increased DNA damage or genomic instability. Examining the transcription factories, as shown in Figs. 2(H-I), we found a slight increase in the cluster size of active RNA polymerase II (phosphorylated Ser5 of the RNAPII C-terminal domain, marking 5’ end of transcribing genes) in normal tissue at risk for tumorigenesis (6-week *Apc*^Min/+^ mice) (*P* < 10^−11^); in tumor cells from 12-week *Apc*^Min/+^ mice, the increase in cluster size and density of active RNAPII becomes substantial (*P* < 10^−15^ and *P* = 0.0005, respectively). This result suggests progressively enhanced formation of active RNAPII clusters during tumorigenesis.

Next, we disrupted heterochromatin structure by knocking down SUV39h1, a histone H3K9 methyltransferase, using small interfering RNA (siRNA), this depletion reduced the total level of H3K9me3 in mouse fibroblast cells and the knockdown efficiency was validated by western blot (Fig. 3E). Super-resolution images of H3K9me3 show that the large and compact heterochromatin foci in control cells disappear upon depletion of SUV39h1, with substantially reduced sizes of H3K9me3 clusters. In contrast, the super-resolution images of DNA (Fig. 3B) reveal that in control cells, the condensed regions of DNA are largely overlapping with H3K9me3 foci; while in cells with SUV39h1 knockdown, the DNA folding becomes more fragmented, as evidenced by the reduced size of nanodomains and the cross-sectional profile (Fig. 3F, H). Analysis of the two-color super-resolution images of DNA/H3K9me3 shows that the percentage of DNA overlapped with H3K9me3 is dramatically reduced upon disruption of heterochromatin structure (Fig. 3H). On the other hand, active RNAPII clusters marked by phosphorylated Ser5 of RNAPII become enlarged with increased level in cells with disrupted heterochromatin (Figs 3C-E, I), indicating enhanced formation of transcription factories. Notably, in control cells, most of the active RNAPII clusters and H3K9me3 nanodomains are spatially exclusive from each other, suggesting that active transcription factories are not located in the heterochromatin domains. Upon disruption of heterochromatin structure in the siSUV39h1 cells, a significantly higher levels of de-compacted heterochromatin domains are co-localized with active RNAPII, as shown in the cross-sectional profiles (Fig 3G, I), suggesting that the spatial juxtaposition between active transcription factories and the de-compacted chromatin domains may promote transcription. Further, we confirmed that cells with SUV39h1 knockdown show an increased number of γ-H2AX foci and chromosomal instability (Fig. 3J-K). Together, these results suggest that disruption of heterochromatin structure leads to fragmented DNA folding, enhanced formation of transcription factories and increased genomic instability.

**Figure 3.**
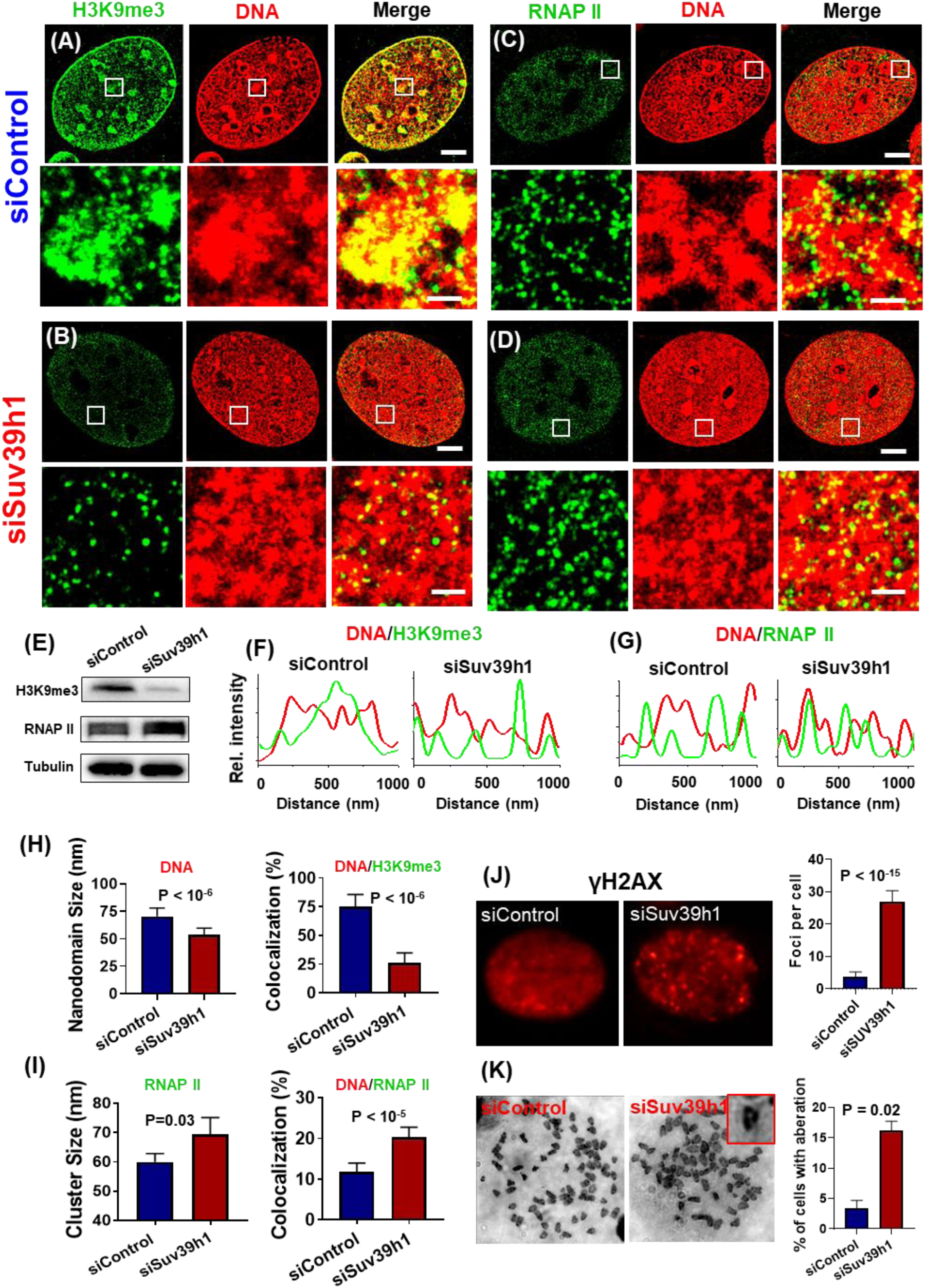
Disruption of H3K9me3 by knocking down SUV39h1 causes decompaction and fragmentation of DNA folding, enhances the formation of transcription factories and induces genome instability. (**A-D**) Representative two-color STORM images showing the spatial relationship between DNA and H3K9me3 (A-B) and between DNA and active (phosphorylated) RNAP II (C-D) in control NIH-3T3 cells and those cells with SUV39h1 knockdown (siSUV39h1). Green: H3K9me3 or phosphorylated RNAP II (labeled by Alexa 647); Red: DNA (labeled by CF568). Scale bars, 4 µm and 500 nm in the original and magnified images, respectively (E) Western blotting of H3K9me3 and phosphorylated RNAP II with tubulin as an internal reference in control cells and those cells with SUV39h1 knockdown (siSUV39h1). (**F-G**) Line profile of signal intensity for DNA and H3K9me3 or RNAP II. (**H-I**) Quantitative analysis of DNA nanodomain size, active RNAP II cluster size and the percentage of DNA that overlaps with H3K9me3 and active RNAP II in control and cells with SUV39h1 knockdown. (**J**) γH2AX immunofluorescence and quantification of γH2AX foci numbers in control and cells with SUV39h1 knockdown. (**K**) Cytogenetic analysis of chromosomal aberration in control cells or SUV39h1 knockdown cells.

### Disrupted higher-order heterochromatin structure is a gradual evolving process and independent of molecular pathways in multiple tumor types

To evaluate whether such structural disruption is due to a specific molecular pathway or a common feature, we imaged heterochromatin structure in another mouse model of intestinal tumorigenesis – *Villin-Cre/BRAF*^V600E/+^ in which the tumor–initiating event in oncogene *BRAF* mutation (mostly V600E) occurs in ∼20% of colorectal carcinogenesis (Rad et al., 2013). We analyzed the cluster size and RDF of H3K9me3-dependent heterochromatin structure from normal cell nuclei of wild-type mice, histologically non-dysplastic cells from *Villin-Cre/BRAF*^V600E/+^ mice at 6 weeks and tumor cells (adenoma) from *Villin-Cre/BRAF*^V600E/+^ mice at 12 months. As shown in Supplementary Fig. S8, the heterochromatin structure shows a substantial decompaction at the early-stage carcinogenesis (6-week *Villin-Cre/BRAF*^V600E/+^ mice) when tissue appears non-dysplastic and this decompaction becomes even more significant in tumor cells (12-month *Villin-Cre/BRAF*^V600E/+^ mice), as reflected in the progressively smaller H3K9me3 cluster size and narrower RDF distribution. This result suggests the disrupted heterochromatin structure in early carcinogenesis is independent of cancer-driven molecular pathways in intestinal tumorigenesis.

Next, we analyzed the *Myc*-driven prostate tumorigenesis mouse model (Hi-MYC), in which prostate cancer is driven by overexpression of Myc under the control of ARR2-probasin promoter (Ellwood-Yen et al., 2003). This model is clinically relevant as *Myc* overexpression was reported in roughly 70% of early-stage prostate cancer (Fox et al., 1993; Jenkins et al., 1997; Qian et al., 1997; Reiter et al., 2000) and share molecular features with the human disease (Ellwood-Yen et al., 2003). As shown in Fig. 4, we analyzed normal tissue from wild-type mice and a set of prostate lesions from Hi-MYC mice with low-grade prostate intraepithelial neoplasia (Low-PIN), high-grade PIN (high-PIN), carcinoma in situ (CIS), and invasive cancer. A similar progressive decompaction of heterochromatin in neoplastic progression of prostate lesions was observed, suggesting a continuous process throughout neoplastic progression (Fig. 4F-G).

**Figure 4.**
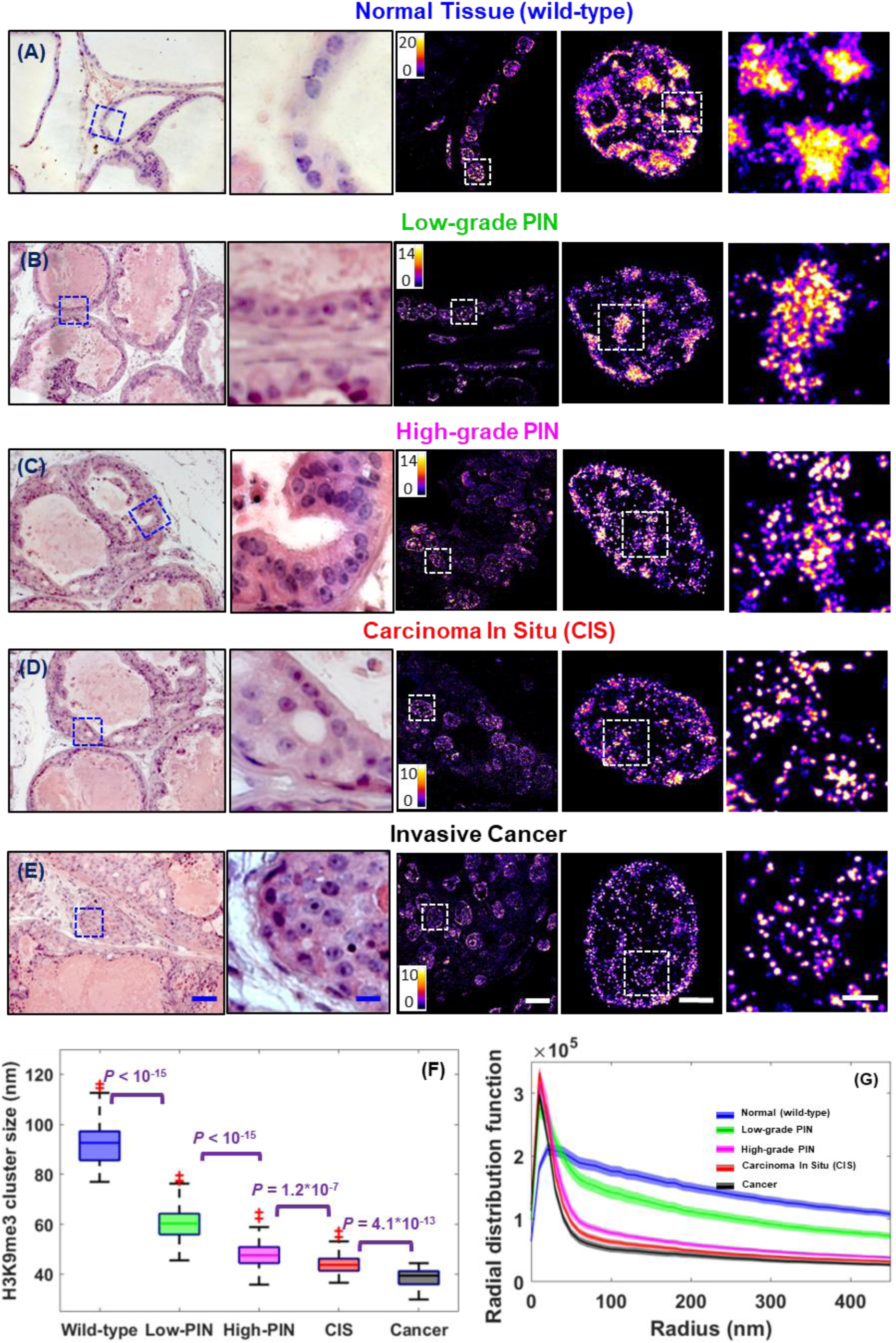
Super-resolution imaging of graudal disruption of the higher-order heterochromatin structure in prostate neoplasia. (**A-E**) Representative histology and the corresponding super-resolution images of heterochromatin structure (from the blue boxes) from normal epithelial cells of the prostate from wild-type mice, low-grade prostate intraepithelial neoplasia (low-grade PIN), high-grade PIN, carcinoma in situ (CIS) and invasive prostate carcionma from Hi-*Myc* mice. Scale bars in the H&E images are 200µm and 10µm, respectively. Scale bars in the STORM images are 10 µm, 2 µm and 500 nm, respectively. (**F**) Box-and-whisker plot of the H3K9me3 cluster size. Over 150 cell nuclei were analyzed for each group. *P* values were determined using Mann-Whitney test. (**G**) Radial distribution function (RDF) that quantifies H3K9me3-dependent heterochromatin structure, averaged over all nuclei for each group. The shaded area shows the standard error.

Further, we used another mouse model that generates pancreatic ductal neoplasm via pancreas-targeted expression of activated oncogene *Kras* (*Kras*^G12D/+^ /*Pdx1-Cre*) and analyzed a spectrum of pancreatic lesions representing various stages of pancreatic neoplastic progression, including normal acinar cells from wild-type mice, acinar to ductal metaplasia (ADM) and various stages of pancreatic intraepithelial neoplasia (PanIN-1, 2, 3). In the *Kras* mouse model, acinar cells are the origin for PanIN lesions, and ADM is the initiating event for the development of pancreatic cancer (Habbe et al., 2008; Reichert et al., 2016; Storz, 2017). As shown in Supplementary Fig. S9, similar to mouse models of intestinal tumorigenesis, we observed a significant decompaction of heterochromatin in the earliest precursor – ADM (Figs. S9 (B1-B4)); the size of heterochromatin nanoclusters undergoes progressive reduction with gradually narrower distribution of RDF during neoplastic transformation (normal acinar cells to ADM, then to PanIN-1, 2 and 3). Therefore, the results shown here using three different tumor types support that progressive decompaction of heterochromatin structure is a common feature across different tumor types throughout neoplastic progression.

### Disrupted higher-order heterochromatin structure is also confirmed in human neoplasia

To confirm our mouse model findings in human tumors, we first analyzed a set of sporadic colorectal neoplasia and their paired normal tissue (see also Supplementary Table S1 for detailed patient characteristics). As shown in Figs. 5(A-D), both STORM images and the RDF show significant size reduction of heterochromatin nanoclusters in human tumors compared to the paired normal tissue. Figure 5E shows the results from 19 patient samples, and a progressive reduction in the size of H3K9me3 nanoclusters was also seen in normal tissue from non-neoplastic patients, adenoma or low-grade dysplasia (LGD), high-grade dysplasia (HGD), and adenocarcinoma (CA) with a high significance. These findings are consistent with our results in the above described mouse models where higher-order heterochromatin structure is progressively disrupted in patient tumors.

**Figure 5.**
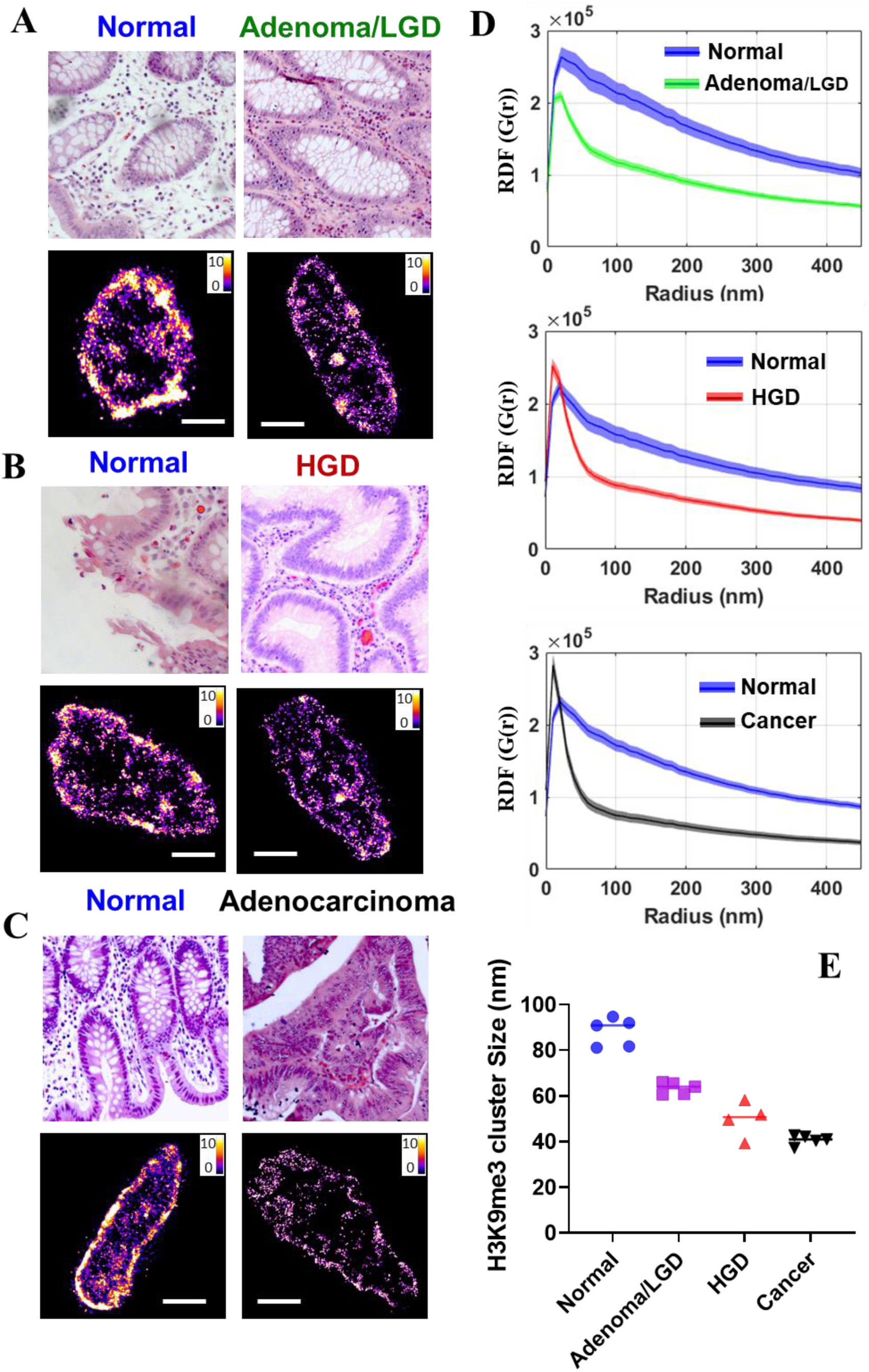
Super-resolution imaging of disrupted higher-order heterochromatin structure in human neoplasia. (**A-C**) Representative histology and corresponding super-resolution images of heterochromatin structure for colorectal neoplastic lesions together with their paired normal tissue located at ∼ more than 10 cm away from the tumor. (**D**) The radial distribution functions (RDF) for characterization of H3K9me3-dependent heterochromatin structure for the normal-tumor pairs. (**E**) The average H3K9me3 cluster size for a total of 19 patient samples (patient informatin is shown in Supplementary Table S1) including 5 normal tissue from non-neoplastic patients, 5 adenoma or low-grade dysplasia (LGD), 4 high-grade dysplasia and 5 invaisve colorectal cancer. Each point is the average of over 150 cell nuclei for each group. *P* values were determined using Mann-Whitney test.

### Disrupted higher-order heterochromatin structure distinguishes precursors that are indistinguishable by conventional pathology

To explore the potential of nanoscale heterochromatin decompaction to detect high-risk precursors that cannot be distinguished by conventional pathology, we generated a more aggressive phenotype with accelerated neoplastic progression in the *Kras*^G12D/+^ mouse model of pancreatic tumorigenesis via inflammatory injury (treating mice with cerulein), which we then compared to the *Kras*^G12D/+^ mice, as well as wild-type mice with and without cerulein treatment. First, we compare the STORM images of normal acinar cells in the pancreas of wild-type mice and *Kras*^G12D/+^ mice (Figs 6A-B). Although acinar cells are normal phenotype, the acinar cells in *Kras*^G12D/+^ mice exhibit heterochromatin decompaction, as evidenced by smaller heterochromatin nanoclusters. Next, the wild-type mice with cerulein treatment developed transient ADM, while *Kras*^G12D/+^ mice developed persistent ADM that eventually progressed into PanIN lesions. As shown in Fig. 6(C-D), we found that the cell nuclei from persistent ADM exhibit more de-compact heterochromatin compared to those from transient ADM. Further, as shown in Fig. 6(E-F), the cell nuclei from PanIN-1 lesion developed from the cerulein*-*treated *Kras*^G12D/+^ mice that underwent accelerated tumorigenesis exhibit more heterochromatin decompaction compared to the same histological entity of PanIN-1 from *Kras*^G12D/+^ mice in the absence of inflammatory insult.

**Figure 6.**
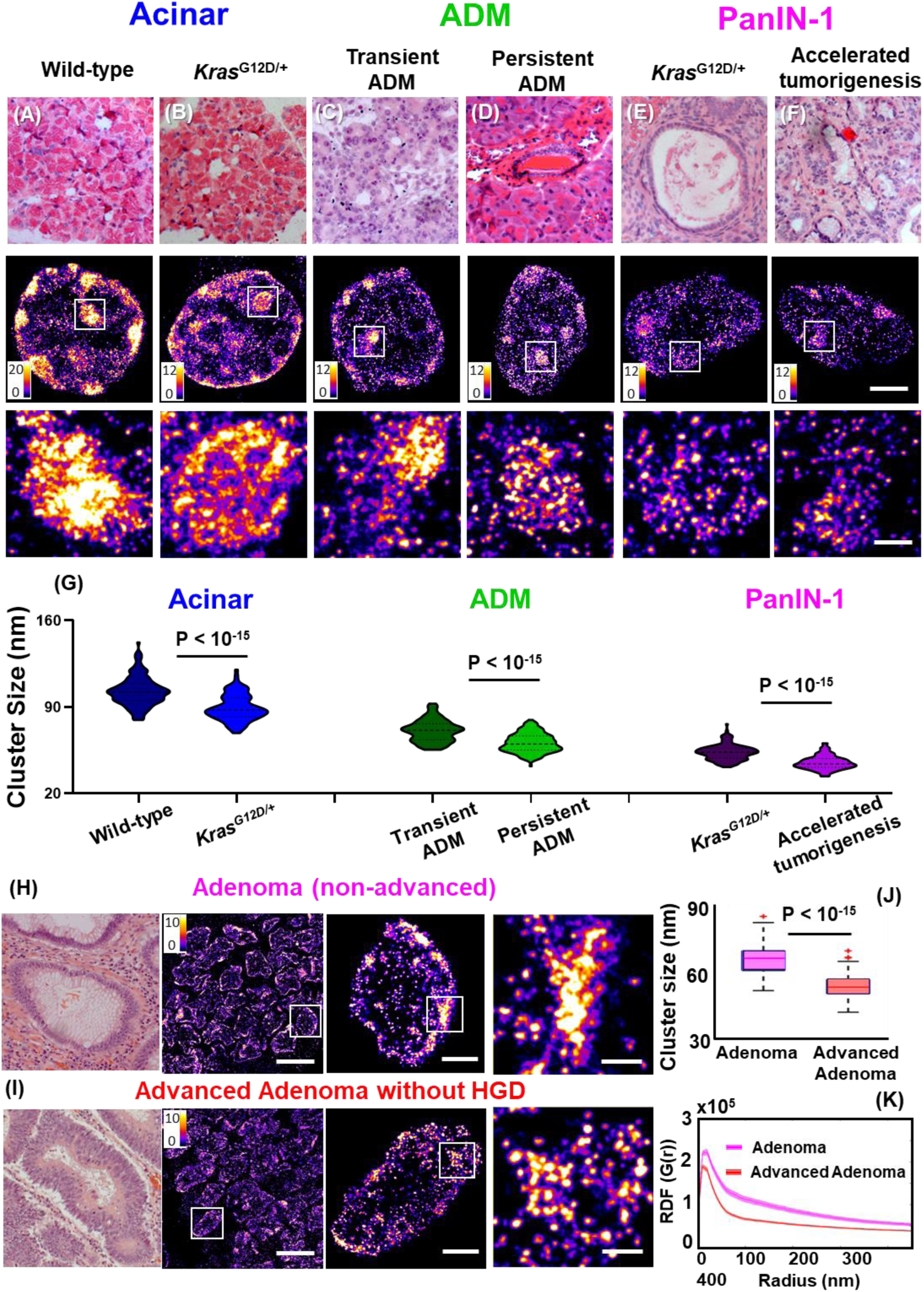
Disrupted higher-order heterochromatin structure by super-resolution imaging distinguishes high-risk precursors that are indistinguishable by conventional pathology. (**A-F**) Representative histology and super-resolution images of heterochromatin structure in normal acinar cells between wild-type mice and *Kras*^G12D/+^ mice, between transient ADM from wild-type mice treated with cerulein and persistent ADM from *Kras*^G12D/+^ mice treated with cerulein, PanIN-1 lesions from *Kras*^G12D/+^ mice and *Kras*^G12D/+^ mice with cerulein treatment that underwent accelerated tumorigenesis. Scale bars in STORM images are 2 µm and 500 nm in the original and magnified images, respectively. (**G**) Statistical analysis of the H3K9me3 cluster size for each group. (**H-I**) Representative histology and corresponding super-resolution images of heterochromatin structure between non-advanced adenoma and advanced adenoma without high-grade dysplasia. Note that adenoma and advanced adenoma without HGD are histologically indistinguisable, and advanced adenoma without HGD refers to those large adenoma with a tumor size of more than 1 cm (1.5 cm polyps). Scale bars in STORM images are 10 µm, 2 µm and 500 nm in the original and magnified images, respectively. (**J**) The box-and-whisker plots of the cluster size of H3K9me3-dependent heterochromatin. Over 150 cell nuclei were analyzed for each group. *P* values were determined using Mann-Whitney test. (**K**) The average radial distribution function (RDF) for H3K9me3, where the solid curve shows the average value from all measured nuclei and the shaded area shows the standard error.

In addition, we analyzed the human precursor lesions that are at higher risk for developing cancer. Figure 6H shows a comparison of the STORM images between non-advanced and advanced adenoma without high-grade dysplasia. These two precursors are histologically indistinguishable, but advanced adenoma presents tumor size of larger than 1 cm that has been clinically established with a higher risk for developing colorectal cancer. The super-resolution images of advanced adenoma indeed present smaller heterochromatin nanoclusters and narrower distribution in RDF, consistent with our findings of more decompacted heterochromatin structure further along neoplastic progression. These results demonstrate that early-stage precursor lesions undergoing more aggressive neoplastic transformation exhibit more de-compact heterochromatin structure, which offers a more sensitive approach to risk-stratify patients at higher risk for aggressive progression.

## DISCUSSION

This work reveals the evolution of higher-order heterochromatin structure and DNA folding throughout all stages of tumorigenesis. These findings were made possible by our optimized STORM-based super-resolution imaging approach and high-fidelity image reconstruction for pathological tissue, referred to as PathSTORM. This study addressed several important unanswered questions in the role of higher-order chromatin folding during tumorigenesis. Super-resolution imaging revealed significant heterochromatin decompaction and fragmented DNA folding at early-stage carcinogenesis prior to the presence of tumor cells (Fig. 1). We cross-validated this finding with an independent biochemical technique, CUT&RUN, which also identified that the most affected genomic regions by the disrupted heterochromatin is satellite repeats. Integrated analysis of CUT&RUN data with RNA-seq also revealed gene misregulation at locations where chromatin structure is altered, and specific upregulation of genes in pathways associated with cancer progression. Disrupting heterochromatin structure leads to increased chromosomal instability, DNA damage, increased formation of active transcription factories and spatial juxtaposition with relaxed chromatin domains. These results provide strong evidence for heterochromatin decompaction as an early event in malignant transformation that results in increased genomic instability and active transcription for cells to gain plasticity that facilitates malignant transformation (Feinberg et al., 2016).

Quantitative analysis of higher-order heterochromatin structure at various stages of carcinogenesis (including normal cells, different stages of precursor lesions and invasive cancer) reveals progressive heterochromatin decompaction, suggesting that such structural disruption is a dynamic and continuous process throughout tumorigenesis, rather than a one-time event. Importantly, we showed that structural disruption in heterochromatin during early carcinogenesis is a common feature independent of molecular pathways in multiple tumor types (colorectal, pancreatic, and prostate). This result further supports the role of heterochromatin decompaction to create an enabling environment that facilitates malignant transformation. Of note, this finding would not be possible without the preserved spatial context of tissue architecture and cell morphology on pathological tissue, which is essential to unambiguously define pathological stages in carcinogenesis, especially for precursor lesions.

Our observed higher-order heterochromatin decompaction is largely in line with previous functional studies of dysregulated heterochromatin in cancer. The loss of heterochromatin proteins or post-translational marks (e.g., H3K9me3, H3K9me2, and HP1) has been widely reported in cancer cells and its functional consequences of impairing genomic stability, changing transcriptional programs, and promoting tumorigenesis and cancer progression (Avgustinova et al., 2018; Carone and Lawrence, 2013; Dialynas et al., 2008; Janssen et al., 2018; Peters et al., 2001; Reddy and Feinberg, 2013; Timp and Feinberg, 2013). While few studies have examined dysregulated heterochromatin in precursor lesions, our study underlines the importance of disrupting the highly compact heterochromatin structure as an initial barrier for malignant transformation. Interestingly, H3K9me3-dependent heterochromatin was shown to be a barrier for cell fate change or reprogramming from somatic cells into induced pluripotent stem cells, which in some extent resembles tumorigenesis (Becker et al., 2016; Feinberg et al., 2016).

Our results of heterochromatin decompaction also provide a molecular-scale explanation for the cytologic feature of chromatin organization in cancer cells that are widely used for cancer diagnosis. Coarse aggregates of heterochromatin was frequently observed in cancer cells under conventional light microscope (Nickerson, 1998; Zink et al., 2004) and even with electron microscope (Cherkezyan et al., 2014). The aggregated heterochromatin may first appear contradictory to our observed structural decompaction and other biochemical studies of heterochromatin loss, but conventional cytology cannot distinguish these two processes. Due to the lack of molecular-scale resolution and quantitative ability, both features appear as enlarged heterochromatin foci under a conventional light microscope. But super-resolution imaging reveals fragmented DNA folding, reduced size of nucleosome nanoclusters, and less DNA molecules at the foci together with the increased occupancy of heterochromatin that collectively suggest structural decompaction rather than aggregation.

Our results suggest a model for the evolution of higher-order heterochromatin structure and DNA folding during malignant transformation from normal to cancer cells, as illustrated in Fig. 7. In normal cells, heterochromatin is highly compacted and spatially segregated from the active transcription factories, preventing the cells from environmental insults. In early carcinogenesis when the cells still appear pathologically normal, the heterochromatin mark (H3K9me3) undergoes a significant reduction, coupled with de-compaction of the dense nucleosome clusters and fragmentation of DNA folding, leading to increased formation of active transcription factories intermingled with DNA regions and genomic instability. As cells transform into tumor cells, the chromatin folding becomes gradually de-compacted and fragmented, with further enhanced formation of active transcription factories. The structure of the heterochromatin foci appears to be enlarged under conventional microscope. Such gradual evolution of heterochromatin decompaction and fragmented DNA folding impair genomic stability and enhance transcription, which set proper environment for cells to gain plasticity to promote malignant transformation.

**Figure 7.**
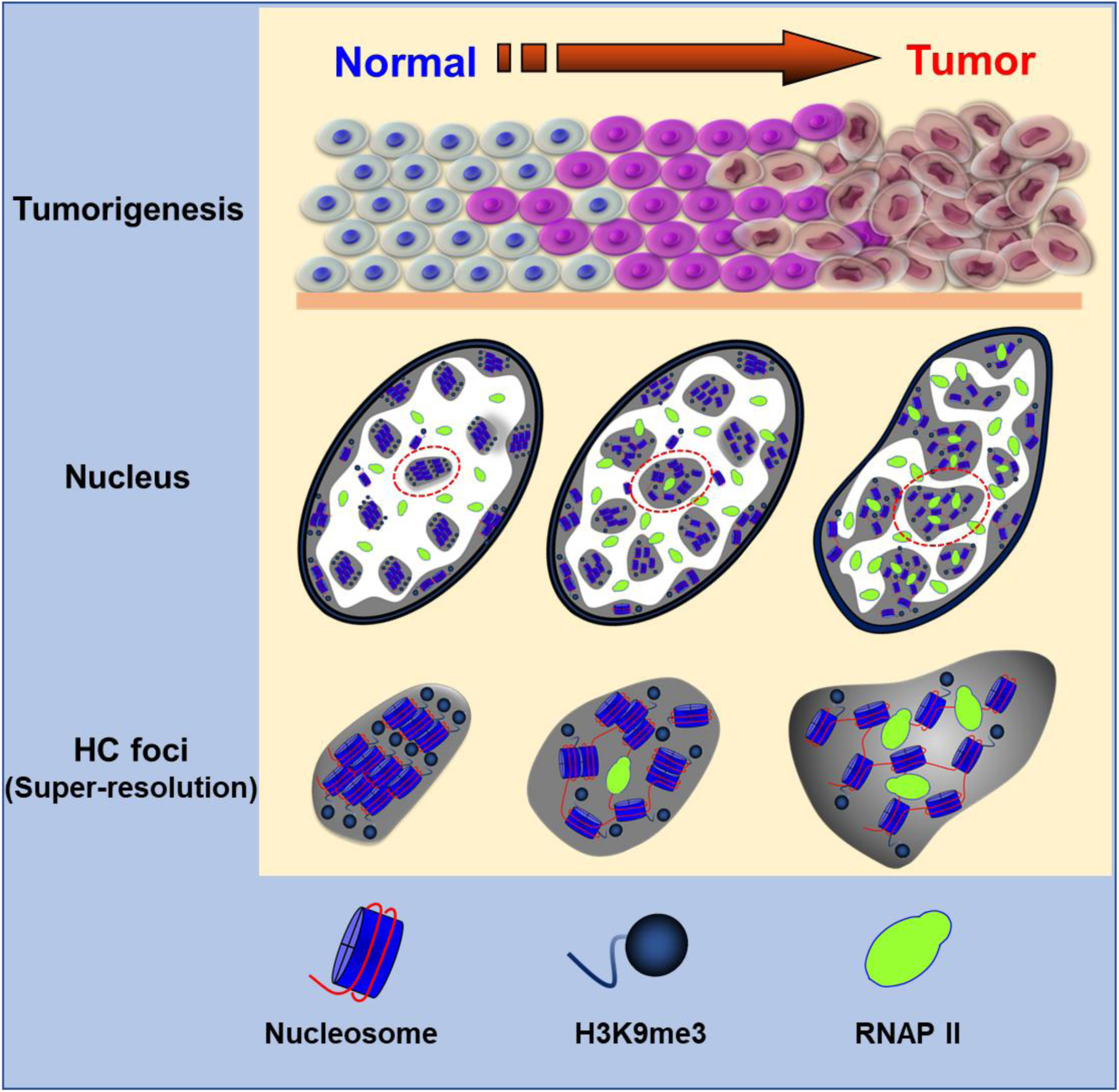
Model to depict the gradual decompaction of higher-order heterochromatin structure, enhanced formation of transcription factories and enlarged heterochromatin foci in carcinogenesis.

Ultimately, new scientific findings need to be translated to improve patient care. We explored the potential of imaging heterochromatin decompaction to improve cancer diagnosis and risk stratification. We showed that in those precursor lesions with the same pathological entity that are histologically indistinguishable, those precursors at higher risk for accelerated malignant transformation present more de-compacted heterochromatin structure in both mouse models of accelerated tumorigenesis and human patients. For example, we showed that in ADM and PanIN-1 precursor lesions of pancreas, those with undergo accelerated tumorigenesis exhibit more de-compacted heterochromatin structure. In adenoma and advanced adenoma without HGD (Fig. 6) that are histologically indistinguishable, the heterochromatin exhibits more severe decompaction in advanced adenoma lesions compared to that from non-advanced adenoma. Although further validation on a large sample size is required, our data demonstrates the possibility of super-resolution imaging of higher-order heterochromatin structure to risk-stratify pre-cancerous lesions.

Taken together, our study reveals nucleosome-level chromatin folding in malignant transformation, which shows gradual, evolving decompaction and fragmented DNA folding as a common feature independent of molecular pathways in multiple tumor types. Our results underline the importance of de-compacted heterochromatin folding to facilitate malignant transformation in early carcinogenesis, which may also be used to improve cancer diagnosis, risk stratification, and facilitate the development and evaluation of new preventive strategies.

## EXPERIMENTAL PROCEDURES

### Immunofluorescence staining

The FFPE tissue blocks taken from mouse models or patients were selected in our study. The 3µm-thick tissue sections mounted on the Poly-D-lysine (PDL)-coated coverslips were deparaffinized in xylene and rehydrated in graded ethanol and finally in distilled water. Cover glasses were then transferred to a pre-heated Tris-EDTA buffer solution (94-96°C, containing 10mM Tris Base, 1mM EDTA, and 0.05% Tween 20, pH 9.0) and heated in a microwave oven for antigen retrieval. Sections were then washed in PBS and then treated with 0.1% solution of sodium borohydride (Sigma-Aldrich) in PBS on ice to reduce autofluorescence. The sections were then permeabilized with 0.2% Trition X-100 (Sigma-Aldrich) in PBS. To block against non-specific binding, the sections were incubated with a blocking solution containing 3% BSA and 0.2% Triton X-100 diluted in PBS for 1 hour at room temperature. Then the sections were incubated with primary antibodies diluted to optimized concentrations at 4°C overnight: rabbit polyclonal to H3K4me3 (1:300, ab8580, Abcam), rabbit polyclonal to H3K27me3 (1:300, Cat.07-449, Millipore), rabbit polyclonal to H4Ac (1:200, Cat.06-598, Millipore), rabbit polyclonal to H3K9me3 (1:300, Cat.07-523, Millipore), mouse monoclonal to RNAP II (1:300), ab5408, abcam), and mouse monoclonal to γ-H2AX (Ser 139) antibody (Santa Cruz Biotechnology, sc-517348). Then Alexa Fluor 647-conjugated donkey-anti-rabbit/mouse secondary antibody was applied to the sections at room temperature for 2 hours in the dark. Sections were then post-fixed in 4% PFA for 10 minutes. The 60% of 2,2-thiodiethanol (TDE, Sigma-Aldrich) in PBS was used to optically clear the tissue sections for ∼20-30 minutes. The fluorescent beads (FluoSpheres™ carboxylate, F8803, Thermo Fisher Scientific) were added into the sample dish as fiduciary markers used for drift correction based on our previously published method (Ma et al., 2017).

### DNA staining in tissue section

In FFPE tissue section, the DNA was stained by TOTO™-3 Iodide (Thermo Fisher Scientific). The tissue section was first treated as described above before the blocking step and then treated with 50 µg/mL DNase-free RNase at 37°C for 30 minutes. Then the tissue section was incubated with 100 nM TOTO-3 for 30 minutes at room temperature. After being washed three times in PBS, the sample was optically cleared as described above, ready for STORM imaging. STORM imaging buffer for TOTO-3 stained tissue consists of 60% TDE (v/v), 10% (w/v) glucose, 0.56 mg/mL glucose oxidase and 0.17 mg/mL catalase.

### STORM image reconstruction and image analysis

The workflow for STORM image reconstruction is shown in Supplementary Fig. S10. Prior to image reconstruction, the background correction based on extreme-value based emitter recovery (Ma et al., 2019) was applied, summarized as follows: (1) transform the intensity value from digital counts to physical photons; (2) segment it into a series of sub-stacks (e.g. 100-frame where the background variation undergoes slow variation); (3) calculate the minimum value of each pixel along the temporal axis for each sub-stack; and (4) calculate the expected background value according to our derived photon statistics model to obtain the expected background value and the temporal minima.

After background correction, we adopted a computationally-efficient emitter-subtraction method, which is a variant of Gaussian deflation method used in high-density localization algorithms (Holden et al., 2011; Sergé et al., 2008) to restore the individual overlapped molecules by subtracting surrounding emitters without the computationally-intensive iterative least squares fitting, followed by our previously developed single-iteration localization algorithm – gradient fitting (Ma et al., 2015) for fast and precise single molecule localization. All above procedures are simple mathematical operation, and thus can be efficiently implemented in GPUs for parallel processing. The reconstructed super-resolution image was rendered by accumulating all the valid molecules with a pixel size of 13 nm.

To perform quantitative analysis of the reconstructed image, we first segmented the epithelial cell nuclei using a semi-automated method described previously (Xu et al., 2018). Subsequent cluster analysis and the radial distribution function (RDF) were performed for each nucleus with a custom software written in MATLAB 2015 (Mathworks) as described previously (Xu et al., 2018). For two-color STORM images, chromatic aberration was corrected by using multi-color fluorescence beads (TetraSpeck microspheres, 0.1-µm diameter, blue/green/orange/dark red fluorescence, Fisher Scientific) as described previously (Xu et al., 2018).

### 3D-SIM image analysis

3D-SIM images were acquired by N-SIM system (Nikon). We performed 3D nucleus segmentation on the reconstructed 3D-SIM image via the following four steps: (1) Background was removed via a 3D median filter with a radius of 5 µm to estimate the local background caused by autofluorescence, which was subtracted from the SIM image. (2) To identify each 3D nucleus along the axial direction, we track the peak intensity distribution along the axial center position of the nucleus. (3) In the initial nucleus segmentation, at the axial position with the maximum intensity, the region of the nucleus was first manually selected by tracking the border of the nucleus, in which the central position was calculated. The precise border of each nucleus was then adaptively defined by finding the fastest changing gradient along each radial direction. This process was repeated for all axial positions above and below the central plane, for 3D nuclei segmentation. Finally, to calculate DNA occupancy, the voxels with an intensity larger than 3 times of square root of the background were recognized as the regions with valid signals. DNA occupancy was defined as the percentage of voxels with valid signals over the total number of voxels for each 3D nucleus.

### CUT&RUN

A total of 4 female mice including 2 *Apc*^Min/+^ mice and 2 age-matched wild-type mice at 6 weeks (Jackson Laboratory) were sacrificed, and their small intestine were removed and washed with PBS. After scraping villi cells from mouse intestine, cells were pelleted, washed with cold PBS, and nuclei were extracted in a hypotonic buffer (NE buffer; 20mM HEPES-KOH pH 7.9, 10mM KCl, 0.5mM Spermidine, 0.1% TritonX-100, 20% glycerol, and freshly added protease inhibitors). CUT&RUN was performed as previously described (Hainer et al 2019). Briefly, lectin-coated Concanavalin A beads (Polysciences) were prepared using binding buffer (20mM HEPES-KOH pH 7.9, 10mM KCl, 1mM CaCl_2_, 1mM MnCl_2_), and nuclei were bound to beads. Bead-bound nuclei were washed (Wash Buffer; 20mM HEPES-KOH pH 7.5, 150mM NaCl, 0.5mM Spermidine, 0.1% BSA, and freshly added protease inhibitors) and primary (H3K9me3, abcam ab8898; H3K4me3, ab8580; or total H3, abcam ab1791) was added to individual reactions. After a 2-hour incubation at 4°C with rotation and two washes in Wash Buffer, recombinant protein A-micrococcal nuclease (MNase) was added to identify the primary antibody. Following two washes in wash buffer, MNase cleavage was initiated with addition of 3mM CaCl_2_, samples were incubated on a water-ice bath for 30 minutes and the reaction was chelated with addition of EDTA and EGTA. RNAs were digested, and protected DNA fragments were released through centrifugation to separate solubilized DNA and protein from insoluble chromatin. Proteins were digested using proteinaseK, samples were purified through phenol chloroform extraction followed by chloroform extraction, and ethanol precipitated with glycogen. Libraries were prepared using Illumina protocols: end repair, A-tailing, and barcoded adapter ligation followed by PCR amplification and size selection. The integrity of the libraries was confirmed by quBit quantification, fragment analyzer size distribution assessment, and Sanger sequencing of ∼10 fragments from each library. Libraries were sequenced using paired-end Illumina sequencing.

To analyze the data, paired-end reads were aligned to mm10 using Bowtie2 (Langmead and Salzberg, 2012) against the entire genome (including repeat regions using RepeatMasker) with the parameters -N 1 and -X 1000. PCR duplicates were removed using Picard (http://broadinstitute.github.io/picard/) and reads with low quality score (MAPQ < 10) were removed using samtools. Reads were separated into 150-500bp for histone modifications. These reads were then be processed in HOMER (Heinz et al., 2010). Mapped reads were aligned over major satellite regions of the genome using the “annotatePeaks” command to make 20bp bins over regions of interest and sum the reads within each bin. Peaks were independently called using the “findPeaks” command, using the “no antibody” control for background signal, to identify all binding sites throughout the genome. Peaks were compared between control and experimental samples using the “mergePeaks” command.

### RNA-seq and transcriptomic analysis

Whole-genome strand-specific RNA-seq was used to profile RNA expression levels in intestinal cells from the same 6-week *Apc*^Min/+^ mice and age-matched wild-type mice used for CUT&RUN experimentation. RNA-Seq libraries were prepared as described previously (Hainer et al., 2015) and in the literature (Kumar et al., 2012). RNA was extracted from intestinal cells using TRIzol followed by column purification (Zymo RNA clean and concentrator column) following the manufacturers’ instructions. Total RNA was depleted of rRNA using a Ribo-Zero Gold kit and first strand cDNA was synthesized. Subsequently, second strand cDNA was synthesized, purified, and fragmented. RNA-seq libraries were prepared using Illumina technology, in a similar way as CUT&RUN libraries described above.

Paired-end reads were aligned to mm10 (mouse models) using RSEM (Li and Dewey, 2011). We assessed the transcripts per million (TPM) changed and visualized the gene expression changes in the wild-type and *Apc*^Min/+^ mice. RSEM output files were used for downstream analysis using HOMER (Heinz et al., 2010). DESeq2 (Love et al., 2014) was used to identify the differentially expressed genes based on significance. To sort the data, K-means clustering was performed using Cluster 3.0 (de Hoon et al., 2004) and heatmaps were generated using Java TreeView (Saldanha, 2004). Gene Ontology (GO) term enrichment was performed to identify gene classes that are mis-regulated using Metascape software (Tripathi et al., 2015). We integrated RNA-seq and CUT&RUN data to determine direct effects of changes in heterochromatin structure on gene expression changes, in which the GO term analysis was performed to identify the direct target molecular pathways due to disrupted heterochromatin structure.

## Statistical methods

The statistical comparison between two groups was calculated using non-parametric Mann-Whitney *U* test in GraphPad Prism 7.0 and two-tailed p-value at 95% confidence interval was presented throughout the manuscript. All the statistical analyses for the tissue samples were based on 150-300 cells. The average cluster size and nearest neighbor distance (nnd) were calculated for each nucleus as a data point. In the box-and-whisker plot, the central line of the box indicates the median; the bottom/top edge of the box indicate 25^th^ /75^th^ percentiles; the whiskers extend to the most extreme data points without outliers.

## Acknowledgments

This work is supported by National Institute of Health Grant Number R01CA185363, R33CA225494 (to Y.L.) and R01CA101753-14 (to S.S.).

