## Supplementary Materials for "Super-resolution imaging reveals the evolution of higher-order chromatin folding in early carcinogenesis"

#### Supplementary Methods

#### Animal models

All animal studies were performed in accordance with the institutional Animal Care and Use Committee of the University of Pittsburgh. All mice were housed in micro isolator cages in a room illuminated from 7:00 AM to 7:00 PM (12:12-hr light-dark cycle), with access to water and chow *ad libitum*.

***$Apc^{Min/+}$ mouse model of intestinal tumorigenesis***

A total of 12 female mice were included in this study for PathSTORM. A first set of three wild-type mice (C57BL/6J, The Jackson Laboratory, Stock No 000664) and three age- and sex-matched *Apc*$^{\text{Min/+}}$ mice (C57BL/6J-*Apc*$^{Min}$/J, The Jackson Laboratory, Stock No 002020) were sacrificed at 6 weeks of age. A second set of three *Apc*$^{\text{Min/+}}$ mice were sacrificed at 12 weeks of age. The 6-week *Apc*$^{\text{Min/+}}$ mice did not show any visible tumor or dysplasia as confirmed by the pathologist; whereas those 12-week *Apc*$^{\text{Min/+}}$ mice had developed multiple visible adenomatous polyps and histologically visible dysplasia in their small intestine. The small intestine tissue was removed, washed with phosphate buffered saline, and prepared in bundles of 1 cm segments. We fixed the tissue in 10% neutral buffered formalin for over 24 hours and embedded the tissue in paraffin block. A segment of small intestine (both proximal and distal parts) was cut, and the tissue was placed in 10% neutral buffered saline for over 24 hours. Then the tissue was embedded in paraffin block.

***Villin-Cre;LSL-BRAF$^{V600E/+}$ mouse model of intestinal tumorigenesis***

The *Villin-Cre;LSL-BRAF*$^{\text{V600E/+}}$ mice (B6.129P2(Cg)-*Braf*$^{tm1Mmcm}$/J, The Jackson Laboratory, Stock No 17837) were generated by crossing *Villin-Cre* mice (B6.Cg-Tg(Vil1-cre)1000Gum/J, The Jackson Laboratory, Stock No 21504) with *LSL-BRAF*$^{\text{V600E/+}}$ mice which were obtained from the Jackson Laboratory (Bar Harbor, ME). Genotyping was performed according to protocols described by the Jackson Laboratory. The *Villin-Cre;LSL-BRAF*$^{\text{V600E/+}}$ mice were euthanized at 6 weeks and 1 year of age. The small intestinal tracts were carefully dissected, rinsed with ice-cold saline, fixed in 10% neutral buffered formalin overnight, and further embedded in paraffin. Small intestine from 6-week-old *Villin-Cre* mice were used as controls.

***Hi-Myc mouse model of prostate tumorigenesis***

Male and female FVB-Tg(ARR2/Pbsn-MYC) mice were procured from the NIH mouse repository (STRAIN 01XK8) and bred in-house following animal protocol approved by the Institutional Animal Care and Use Committee of the University of Pittsburgh. After genetic verification by PCR, 3 pairs of mice (3 wild-type and 3 Hi-Myc mice) at 5 weeks of age were fed with AIN-93G diet and sacrificed at 26 weeks of age. Prostate tissues were collected at the time of sacrifice and fixed in 10% neutral buffered formalin and paraffin-embedded.

***Pdx1-Cre;LSL-KRAS$^{G12D/+}$* mouse model of pancreatic tumorigenesis**

The *Pdx-Cre* (B6.FVB-Tg(Pdx1-cre)6Tuv/Nci, STRAIN 01XL5) and *LSL-KRAS*$^{G12D/+}$ (B6.129-*Kras*$^{tm4Tyj}$/Nci, STRAIN 01XJ6) mice were received from the NCI Mouse Repository. These mice were crossed to generate *Pdx-Cre;LSL-KRAS*$^{G12D/+}$ mice. The genotyping was performed according to the protocols described by the NCI Mouse Repository. The three *Pdx-Cre;LSL-KRAS*$^{G12D/+}$ mice at the age of 7-month were euthanized, the pancreas was dissected, fixed overnight in 10% neutral buffered formalin, and embedded in paraffin. STORM imaging was performed in the pancreatic intraepithelial neoplasia (PanIN) lesions graded at PanIN-1, PanIN-2 and PanIN-3. In addition, we treated three 6-week-old C57BL/6 mice (Jackson Laboratory, Bar Harbor, ME) and three age-matched *KRAS*$^{G12D/+}$ mice by six hourly intraperitoneal injection of caerulein (Sigma-Aldrich) dissolved in PBS on two consecutive days at a dose of 50 µg/kg. Pancreas from wild-type and *KRAS*$^{G12D/+}$ mice were harvested at two days and 21 days, respectively, after the last injection. The normal acinar cells were imaged from the pancreatic tissue of three C57BL/6 mice at 6-7 weeks. After the pancreatic tissue was harvest, it was fixed overnight in 10% neutral buffered formalin and embedded in paraffin.

**STORM setup**

STORM images were acquired using our custom-built system on the Olympus IX71 inverted microscope frame with a 100x, NA=1.4 oil immersion objective (UPLSAPO 100XO; Olympus) and the system has been described in detail previously (Ma et al., 2017; Xu et al., 2018). For single-color dSTORM imaging, the excitation laser at 642 nm (VFL-P-1000-642-OEM3; MPB Communications, Point-Claire, Quebec, Canada) was used at power density of ~2.5kW·cm$^{-2}$ for STORM imaging. The exposure time was 20 milliseconds and a total frame number of 40,000 were used. During the image acquisition, a small amount of activation power (~1µW) for 405 nm laser (DL405-050, CrystaLaser, Reno, NV) was added at 3001$^{st}$ frame and the power of 405 nm laser was gradually increased at a rate of 0.2% per 1000 frames. Two-color dSTORM imaging was conducted sequentially, where first 30,000 frames were acquired on Alexa Fluor 647 with an exposure time of 20 ms for each frame, followed by 30,000 frames of Cy3B with the same exposure time. The excitation laser of 561 nm at laser power of 0.8kW·cm$^{-2}$ (VFL-P-200-560-OEM1, MPB Communications, Point-Claire, Quebec, Canada) was used for imaging Cy3B-labeled targets. The drift correction was independently performed every 200 frames (or 4 seconds) with fluorescent beads (Thermo Fisher Scientific, F8803) excited with 488 nm laser (DL488-150, CrystaLaser, Reno, NV) as fiduciary markers throughout each image acquisition process, based on our established method (Ma et al., 2017). The imaging conditions (exposure time, power density, activation, frame number) remain the same for all experiments reported in this study.

**STORM imaging buffer**

STORM imaging buffer for cultured cells contains 10% (w/v) glucose (Sigma-Aldrich), 0.56 mg/mL glucose oxidase (Sigma-Aldrich), 0.17 mg/mL catalase (Sigma-Aldrich), 0.14M 2-mercaptoethanol (βME, Sigma-Aldrich). For FFPE tissue section, to reduce the high background caused by the strong scattering of pathological tissue and match the refractive index, optical clearing process were conducted before imaging by immersing the sample in 60% (v/v) 2,2'-thiodiethanol (TDE) for 20-30 minutes to make the sample transparent. For the STORM imaging buffer of FFPE tissue section, 60 % (v/v) TDE solution was used instead of water to match the tissue's index, and contains 10% (w/v) glucose (Sigma-Aldrich), 0.56 mg/mL glucose oxidase (Sigma-Aldrich), 0.17 mg/mL catalase (Sigma-Aldrich), 0.14M 2-mercaptoethanol (βME, Sigma-Aldrich), and 0.2 mM Cyclooctatetraene (COT, Sigma-Aldrich). The STORM imaging buffer for ultrathin tissue section contains the same reagents at the same concentration, but without TDE. The imaging buffer for TOTO-3 labeled DNA in tissue contains the same reagents at the same concentration, but without COT and βME. The imaging buffer was added into the sample dish right before imaging.

**Western blotting**

Western blot analysis was performed following the standard protocols with tubulin as loading control. In brief, cell lysates were resolved in SDS loading buffer and subjected to electrophoresis in SDS-polyacrylamide gels and transferred to the polyvinylidene difluoride (PVDF) membrane. After blocking with 5% non-fat milk for 1 hour, membranes were incubated with corresponding primary antibody (H3K9me3, abcam, 1:400; tubulin, Cell Signaling Technology, 1:3000; RNAP II, abcam, 1:1000) at 4°C overnight. The membranes were washed three times in PBS and incubated with horseradish peroxidase (HRP)-conjugated secondary antibody (abcam, 1:2000) for 1 hour at room temperature. Membranes were washed 3 times with 0.1% PBS before exposure. Detection was done using BIORAD Universal Hood II machine with ImageLab software.

**Metaphase spreading**

NIH3T3 cells reaching 70-80% confluency were treated with 0.1 μg/ml Colcemid (Thermo Fisher Scientific) for 1 hour, then trypsinized and washed with PBS. After centrifugation, cells were resuspended with 75 mM KCl for 20 minutes at 37 °C. Cells were fixed with freshly made Carnoy's Fixative (3+1 v/v methanol/glacial acetic acid) three times. Cells suspension was dropped on an ice-cold clean slide and air-dried for one day. Slides were stained with KaryoMAX™ Giemsa Stain Solution (Thermo Fisher Scientific) and observed under the bright-field microscope.

**Sample Preparation for 3D-SIM imaging**

Three C57BL/6J wild-type mouse and three age-matched *Apc*$^{Min/+}$ mice were sacrificed and their small intestine tissue were removed, fixed in 2% paraformaldehyde solution in PBS and placed in 30% sucrose for cryoprotection. The tissue was then flash frozen in liquid nitrogen and then stored at -80°C. Prior to imaging, the frozen tissue was sectioned at 15 µm using a cryostat and stained with 4′,6-diamidino-2-phenylindole (DAPI) and mounted in Gelvatol. A 3D stack of fluorescence images were acquired using N-SIM (Nikon) with a scanning depth of 18 µm at a step size of 0.12 µm. The 3D-SIM images were reconstructed using the image reconstruction software on the N-SIM system.

**Ultrathin tissue section**

A C57BL/6J wild-type mouse was sacrificed and the small intestine tissue was removed, washed with PBS. Two adjacent equal segments were taken. One piece was fixed in 10% formalin for 2 hours and then processed using standard paraffin-embedding protocol and sectioned at 3 µm. The immunofluorescence staining and PathSTORM imaging were performed on the FFPE tissue section as described in the Experimental Procedures of the main text. The second piece was fixed in 2% paraformaldehyde for 2 hours, then in 30% sucrose for 24 hours, and stored in liquid nitrogen. Then an ultrathin section (~700 nm) was cut using an Ultramicrotome (Reichert Ultracut). The standard immunofluorescence staining was performed and STORM imaging was done in the same way as previously described in cultured cells (Xu et al., 2018).

**DNA and immunofluorescence staining for cultured cells**

The DNA staining in cell cultures was performed by using the Click-iT Plus EdU (5-ethynyl-2′-deoxyuridine) Alexa Fluor Imaging Kit (Thermo Fisher Scientific), as described in detail previously (Xu and Liu, 2019; Xu et al., 2018). In brief, cells were incubated with 1 µM EdU contained medium for 24 hours, after fixation and permeabilization, cells were blocked with 3% BSA and incubated with EdU Click-iT Plus reaction cocktail for 30 minutes following the manufacturer’s instruction. DNA were detected by Azide CF-568. For two-color staining of DNA and proteins, after being washed out of the reaction cocktail, cells were incubated with primary antibody at 4 °C overnight. The cells were then washed 3 times with the washing buffer for 5 minutes per wash, and the corresponding Alexa-647 conjugated secondary antibodies were added to the sample in blocking buffer and incubated for 2 hr at room temperature. The cells were washed again 3 times with washing buffer and once with PBS for 5 min per wash and stored in PBS before imaging.

**DNA staining at different stages of cell cycle**

Cells at ~ 60% confluency were incubated with 1 µM EdU for 24 hours, then the medium was changed to the fresh medium containing 10 µM BrdU (Thermo Fisher Scientific) and the cells were incubated for another 1 hour. Cells were fixed with 4% PFA for 15 minutes and permeabilized with 0.2% Triton X-100 for 10 minutes. DNA was denatured by incubating with 2N HCl for 30 minutes at 37°C to make the nucleotides accessible for the antibody, then neutralized by incubating with Tris-HCl buffer pH 7.5 for 20 minutes. The cells were blocked in 3% BSA for 2 hours and incubated with BrdU antibody (Cell

Signaling Technology) at 4°C overnight. After washing, cells were incubated with Cy3B-conjugated secondary antibody for 2 hours at room temperature. The cells were again washed 3 times, and the EdU was detected by Click-iT Plus EdU Alexa Fluor Imaging Kit as descried above. Azide Alexa Fluor 647 was then used to detect EdU labeled DNA.

**RNA interference**

NIH3T3 cells were cultured in DMEM with 10% FBS at 37 °C and 5% $CO_2$. Before transfection, cells were plated on 2 cm MatTek dish until they reach 70% confluency. Cells were transfected with 50 nM Suv39h1 siRNA (Integrated DNA Technologies) using RNAimax transfection reagent (Thermo Fisher Scientific) diluted in OptiMEM (Gibco), after 24 hours, medium was exchanged to fresh medium with 1 μM EdU and incubated for another 24 hours. Cells were fixed with 4% PFA for 15 minutes and permeabilized with 0.2% Triton X-100 for 10 minutes. Cells were stored in PBS for DNA or immunofluorescence staining.

**Hematoxylin & Eosin (H&E) Staining**

FFPE sections were deparaffinized in xylene and rehydrated in graded ethanol/water followed by distilled water. Then the sections were stained with Harris hematoxylin (Anatech Ltd), washed by distilled water, then treated with 10% glacial acetic acid (Fisher Scientific) and Scott’s Tap Water Substitute (1 min, Cancer diagnostics Inc). Then the tissue was stained with Eosin-Y (Anatech Ltd), followed by three washes with 70% ethanol, three washes with 100% ethanol and three washes with xylene. Finally, stained sections were cover slipped with microscope cover glasses (Fisherbrand, Fisher Scientific) and dried at room temperature.

### Supplementary Figures

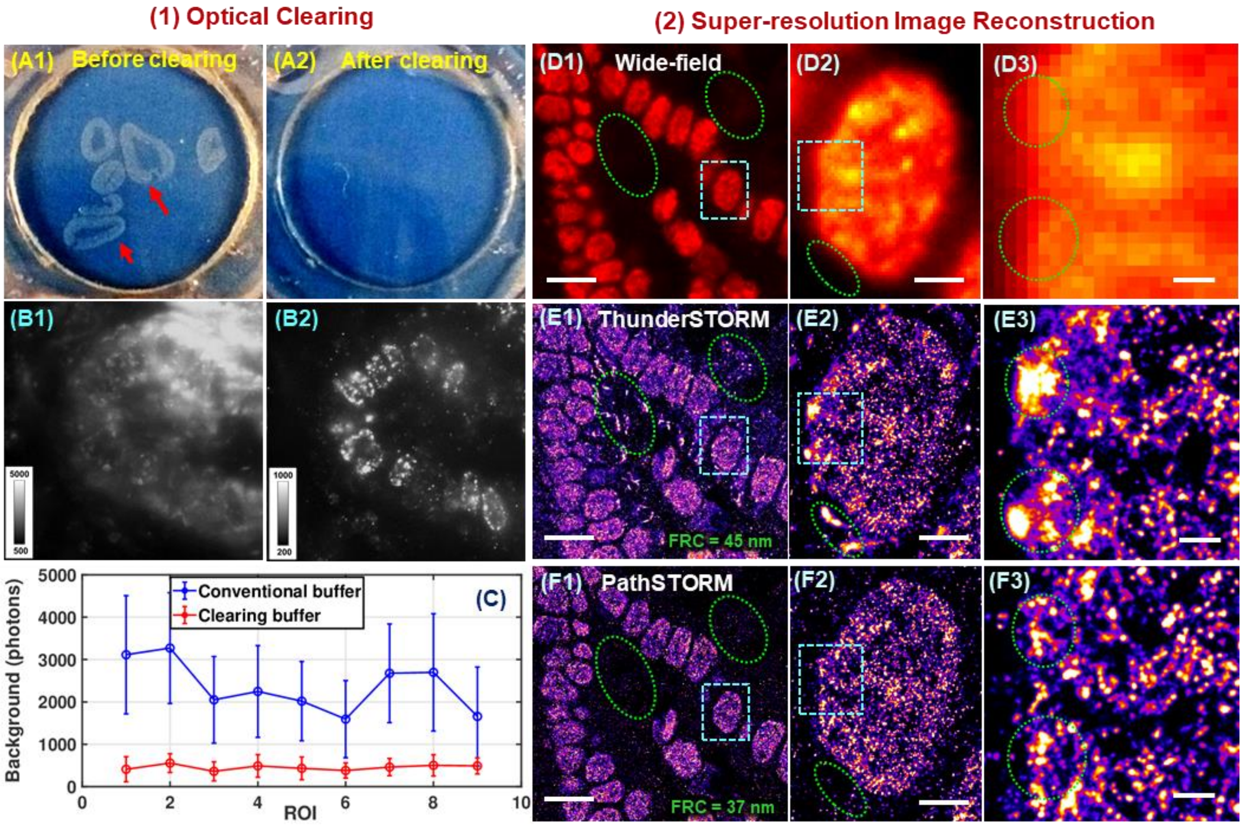


**Figure S1.** The workflow of PathSTORM. (1) Optical clearing: **(A1)** Mouse intestinal tissue section (3µm) before optical clearing and **(B1)** the raw image in the standard aqueous-based STORM imaging buffer at a power density of ~3kW/cm$^2$. **(A2)** After optical clearing, tissue appears transparent and **(B2)** the raw image shows dramatically reduced background under index-matched imaging buffer. **(C)** The comparison of average background signals before and after clearing from 9 different regions of interest (ROIs). The error bar is the standard deviation of background signal for each ROI. There is about 5-fold reduction in background after clearing. (2) Super-resolution image (H4Ac) reconstruction that corrects for heterogeneous background and decomposes overlapping emitters. **(D-F)** The conventional wide-field image **(D1-D3)** and the reconstructed super-resolution images with **(E1-E3)** a conventional method (ThunderSTORM) and **(F1-F3)** PathSTORM. **(D2-D3)**, **(E2-E3)** and **(F2-F3)** are the progressively zoomed images of **(D1-F1)** in the blue square, respectively. The scale bars in (D1-F1), (D2-F2) and (D3-F3) represent 10 µm, 2µm and 500 nm, respectively. The Fourier Ring Resolution (FRC) resolution is shown at the bottom of (E1-F1). Comparing the same areas in the conventional diffraction-limited wide-field images and the reconstructed STORM images by ThunderSTORM and PathSTORM as shown in (D-F), the image reconstructed by conventional method ThunderSTORM shows apparent image artifacts that are not present in the diffraction-limited image. In the zoomed super-resolution image (E2-E3), the artifacts (circled in green) distort the image of the actual chromatin structure. In contrast, the image reconstructed by PathSTORM (F1-F3) shows good agreement with the diffraction-limited image with a higher image resolution. These results showed that PathSTORM increases the image resolution and reduces the image artifacts present in chromatin structure and background.

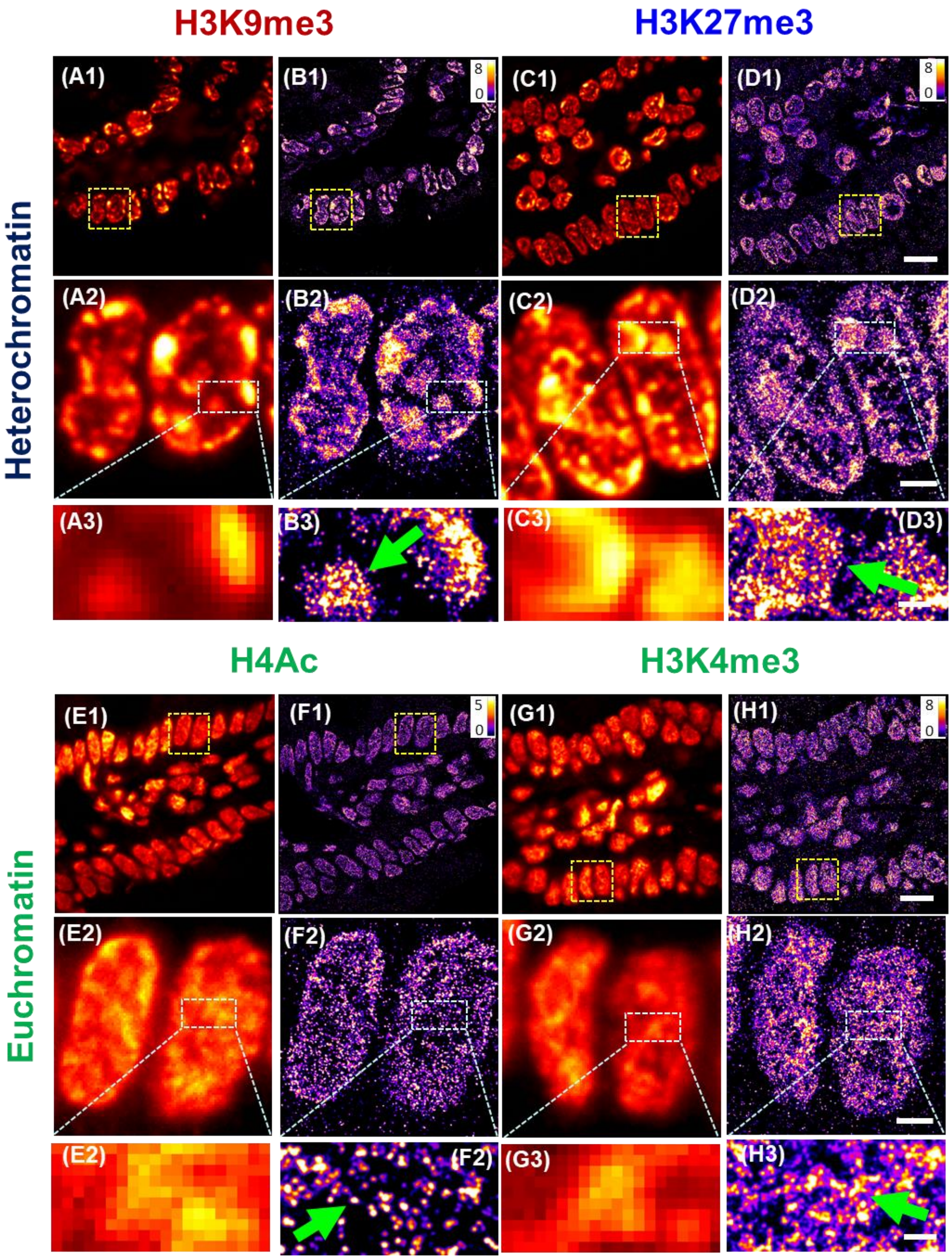


**Figure S2.** Representative (**A1**, **C1**, **E1**, **G1**) conventional wide-field and (**B1**, **D1**, **F1**, **H1**) super-resolution images of (**A1-D1**) higher-order heterochromatin structures marked by transcriptionally repressive histone proteins (H3K27me3 and H3K9me3) and (**E1-H1**) higher-order euchromatin structures marked by transcriptionally active histone marks (H4Ac and H3K4me3) on the pathological tissue of mouse intestine. (**A2-H2**) and (**A3-H3**) are the progressively zoomed images of (A1-H1). The scale bars of (A1-H1), (A2-H2) and (A3-H2) represent 10 µm, 2 µm and 500 nm, respectively. The green arrows indicate the characteristic condensed large nanoclusters formed by heterochromatin and more uniform or spatially diffuse nanoclusters formed by euchromatin.

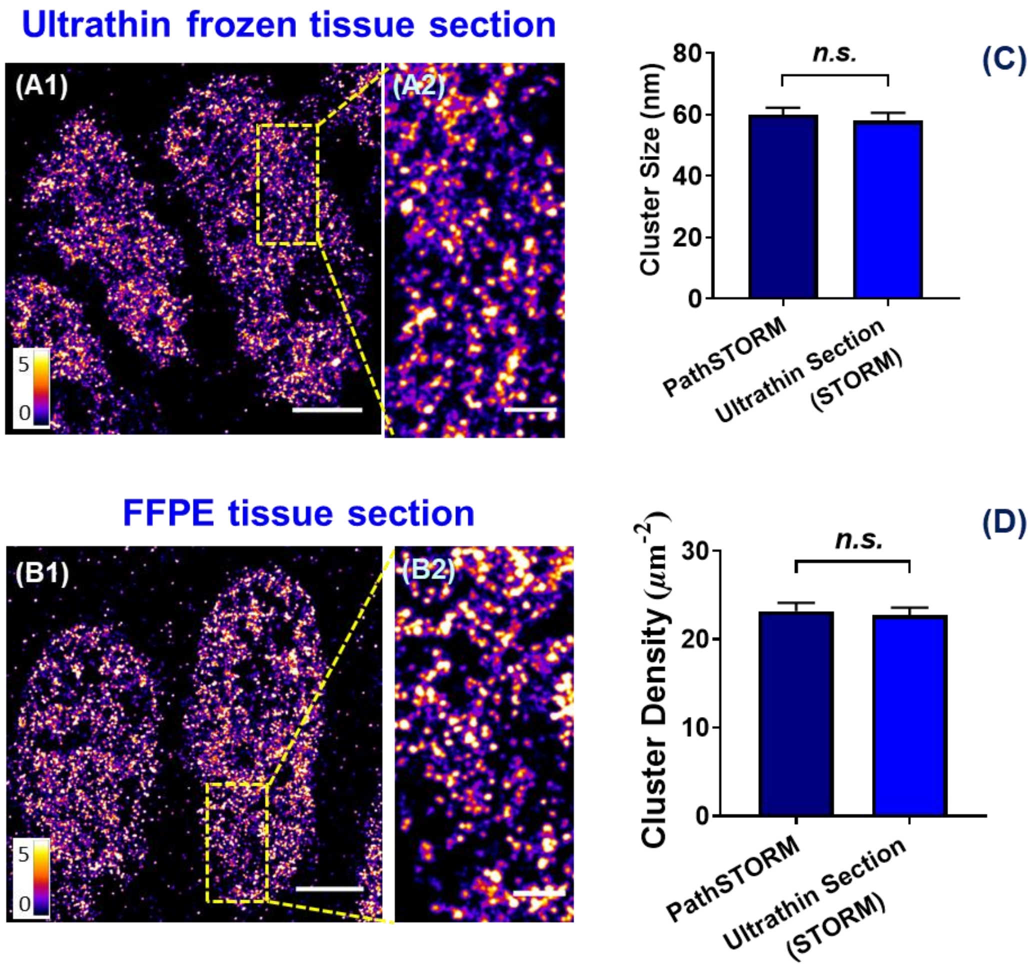


**Figure S3. (A1-B1)** Comparison of the reconstructed STORM images of euchromatin structure (stained with euchromatin marker H3K4me3) from the ultrathin frozen section (**A1-A2**) and FFPE section (**B1-B2**) of the same mouse small intestine stained with H3K4me3. The scale bar in (A1, B1) and (A2, B2) is 2 µm and 500 nm, respectively. (**C-D**) Comparison of euchromatin cluster size (C) and cluster density (D) for the tissue from two closely adjacent areas with one processed as FFPE tissue section and imaged with PathSTORM; another processed as ultrathin frozen tissue section and imaged with standard STORM imaging. The ultrathin frozen section (700 nm) was cut with ultra-microtome, stained and imaged with standard protocols used for cultured cells. The FFPE tissue section was cut at 3µm with a microtome, stained, imaged and reconstructed with PathSTORM method described in the main text.

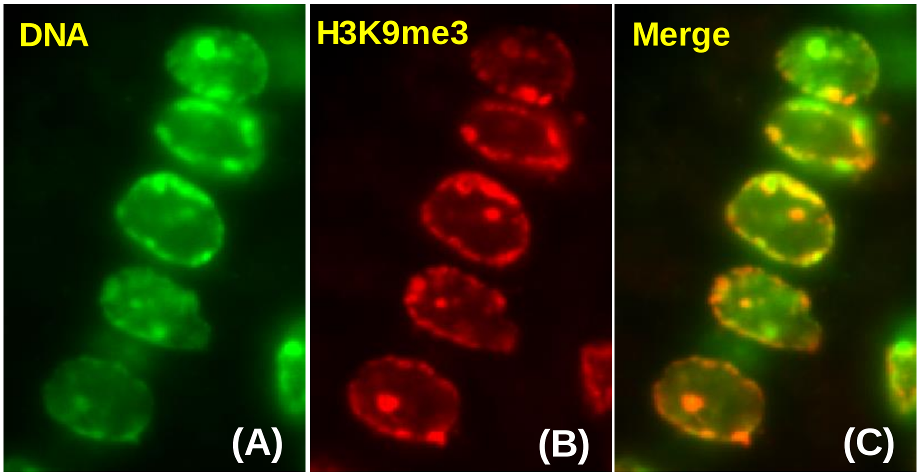


**Figure S4.** Wide-field fluorescence images of (**A**) DNA (stained with DAPI) and (**B**) H3K9me3 (Cy3B) and (**C**) the merged image on mouse intestinal epithelial tissue. Evidently, the histone mark H3K9me3 largely overlaps with the condensed regions of the DNA in the cell nuclei *in vivo*.

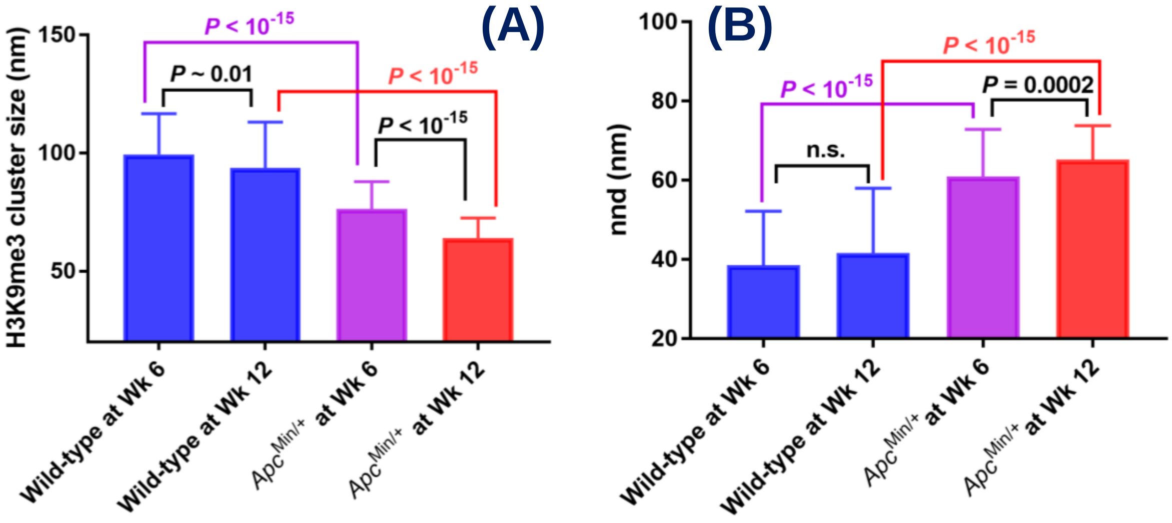


**Figure S5.** The box-and-whisker plots of (A) H3K9me3 cluster size and (B) nearest neighbor distance (nnd) from intestinal epithelial cell nuclei of normal tissue from wild-type mice at 6 weeks and 12 weeks, compared to the age-matched *Apc*$^{Min/+}$ mice. The cluster size of H3K9me3 only shows marginal difference, and nnd shows no difference between 6-week (n = 204 normal epithelial nuclei analyzed) and 12-week (n = 241 normal epithelial nuclei) wild-type mice. The age-related difference is significantly smaller compared to carcinogenesis-associated difference.

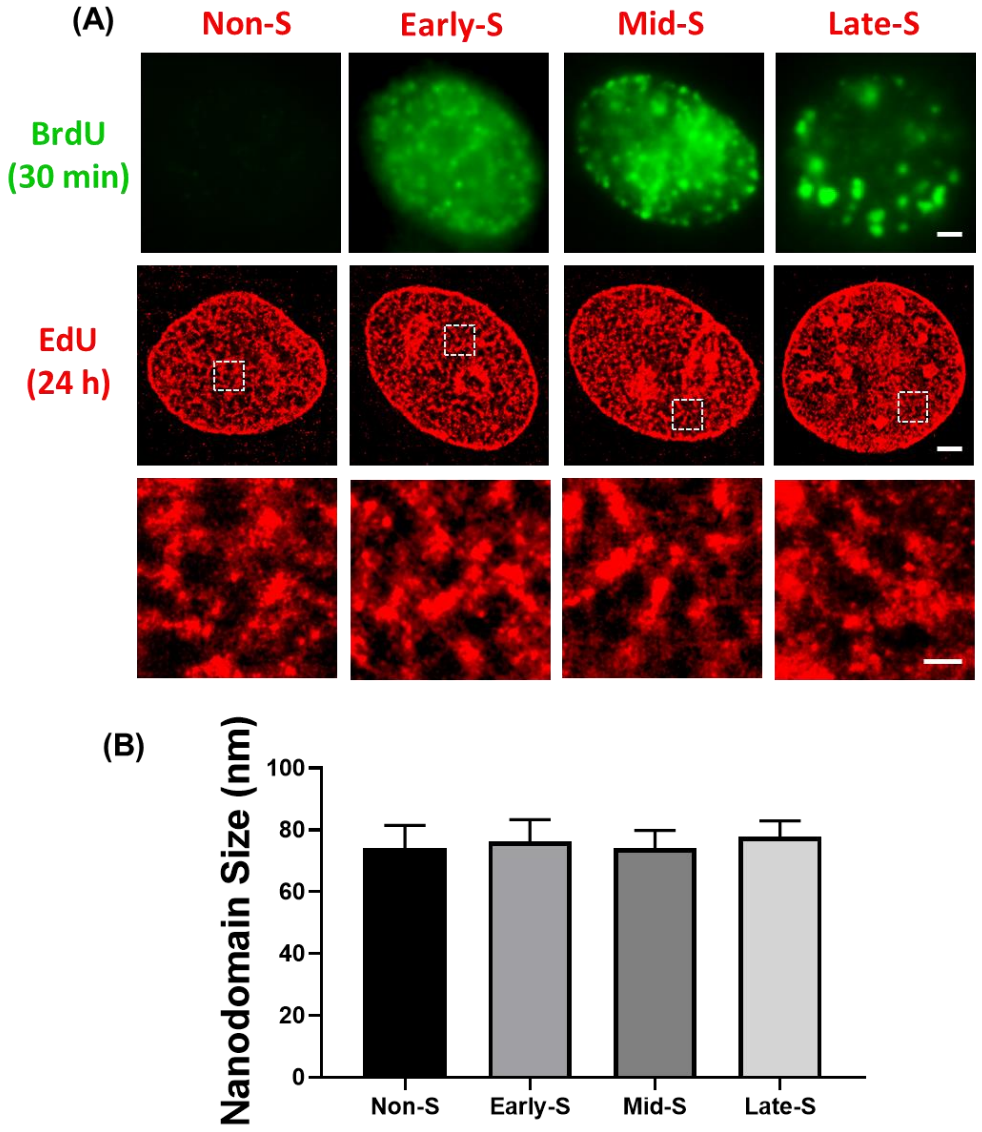


**Figure S6. (A)** Super-resolution images of DNA structure at different cell cycles. BrdU is used to mark cells at S-phase; EdU is used to stain genome-wide DNA. The scale bars represent 2 µm, 2 µm, and 500 nm, respectively. **(B)** Statistical analysis of DNA nanodomain size of different cell cycles. There is no statistical significance in the average size of the nanodomains.

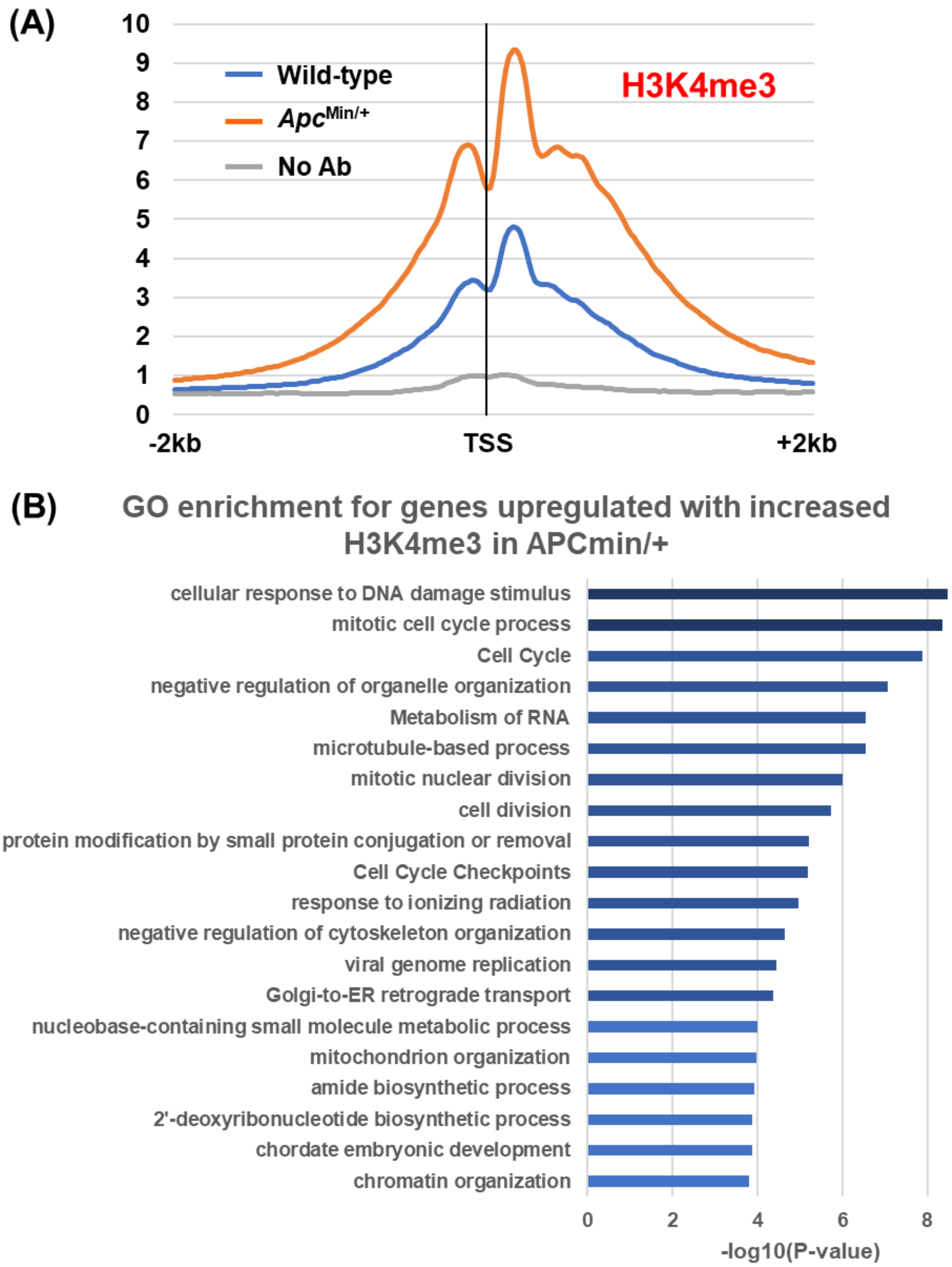


**Figure S7. (A)** The average enrichment of H3K4me3 at transcription start sites (TSS) from epithelial tissue from 6-week wild-type mice (blue), 6-week *Apc*$^{Min/+}$ mice (orange), and no primary antibody control (gray). **(B)** The gene ontology (GO) analysis was performed using Metascape software (Tripathi et al., 2015) for overlapping up-regulated genes with increased occupancy of H3K4me3 in *Apc*$^{Min/+}$ mice.

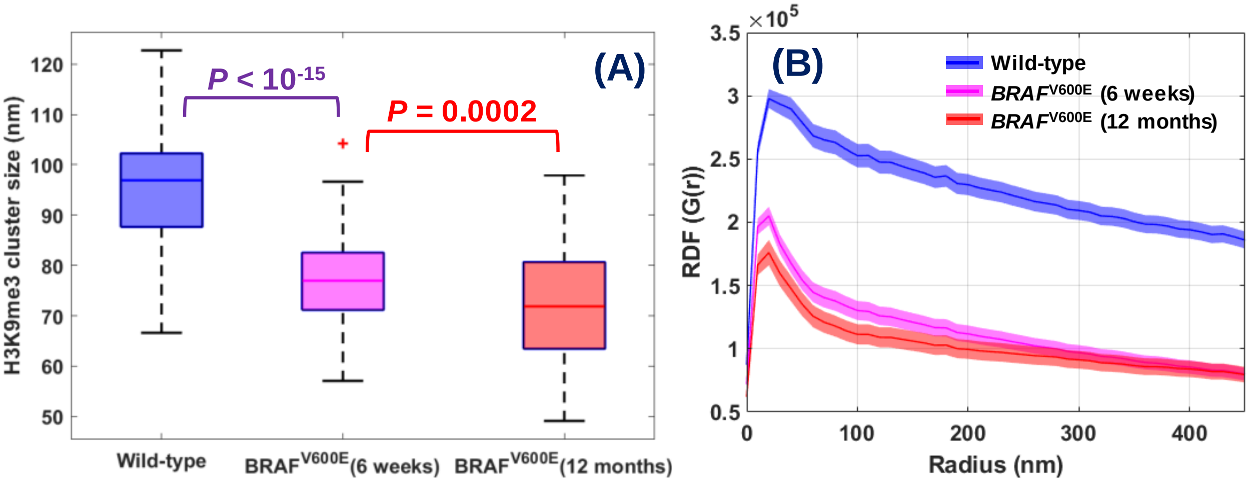


**Figure S8. (A)** The box-and-whisker plots of the H3K9me3 cluster size from intestinal epithelial cell nuclei of normal tissue from wild-type mice, histologically normal-appearing tissue from 6-week *BRAF*$^{V600E}$ mice and tumor (adenoma) from 12-month *BRAF*$^{V600E}$ mice. About 100-200 epithelial cell nuclei were analyzed for each group. *P* values were determined using Mann-Whitney test. (**B**) Average radial distribution function (RDF) for all nuclei in each group. The solid curve shows the average RDF from all measured nuclei and the shaded area shows the standard error.

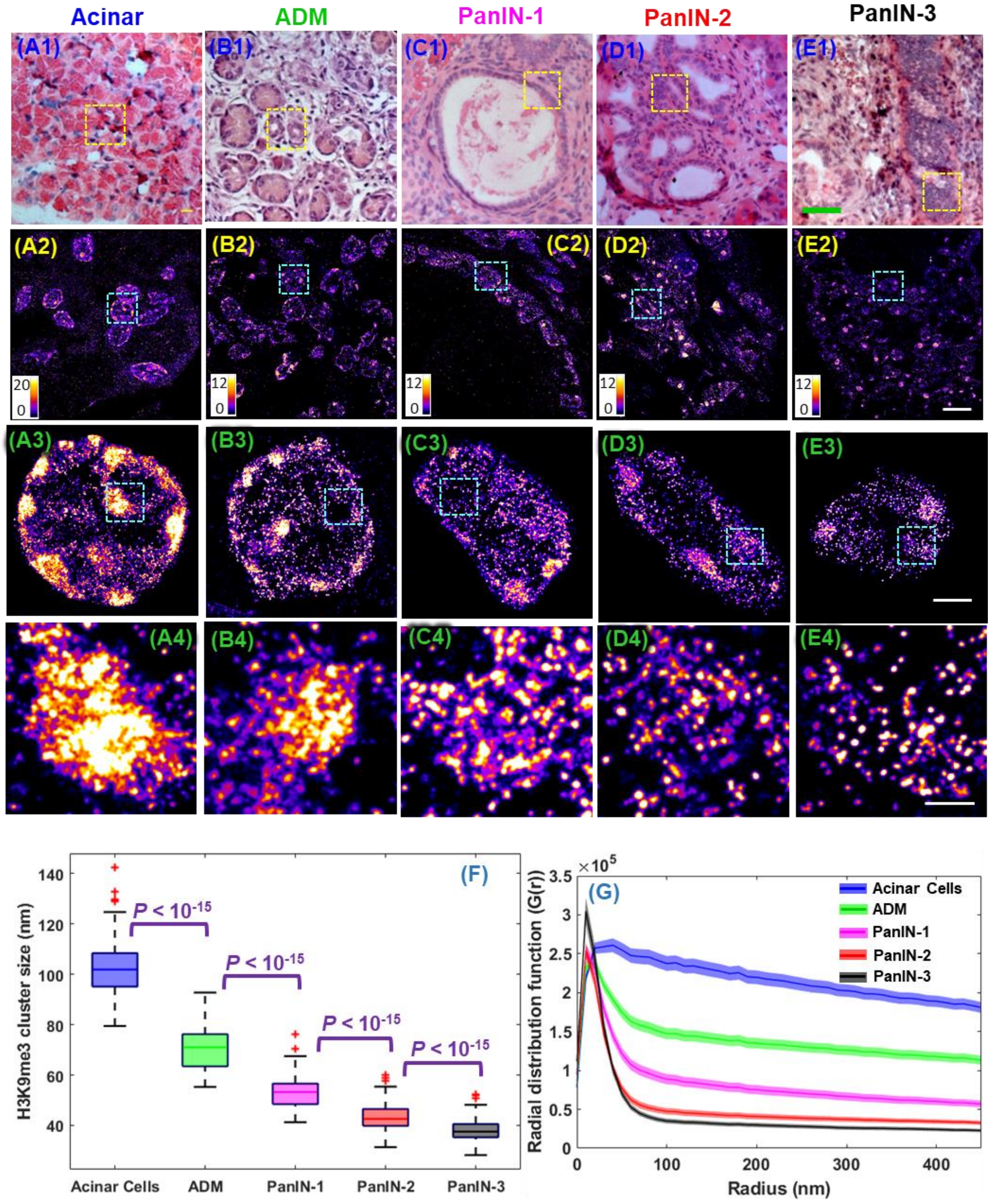


**Figure S9.** Representative (**A1-E1**) histology (scale bar: 200 µm) and (**A2-E2**) super-resolution images of H3K9me3-dependent heterochromatin structure from normal acinar cells of the pancreas from wild-type mice, acinar-to-ductal metaplasia (ADM), pancreatic intraepithelial lesions (PanIN) grade 1, 2 and 3 from *Pdx1-Cre KRAS*$^{G12D/+}$ mice. (**A3-E3**) and (**A4-E4**) are the progressively zoomed images of (A2-E2). The scale bars of (A2-E2), (A3-E3) and (A4-E4) represent 10µm, 2µm and 500 nm, respectively. (**B**) Box-and-whisker plot of the H3K9me3 cluster size. (**C**) Radial distribution function (RDF) that quantifies H3K9me3-dependent heterochromatin structure, averaged over all nuclei for each group. The shaded area shows the standard error.

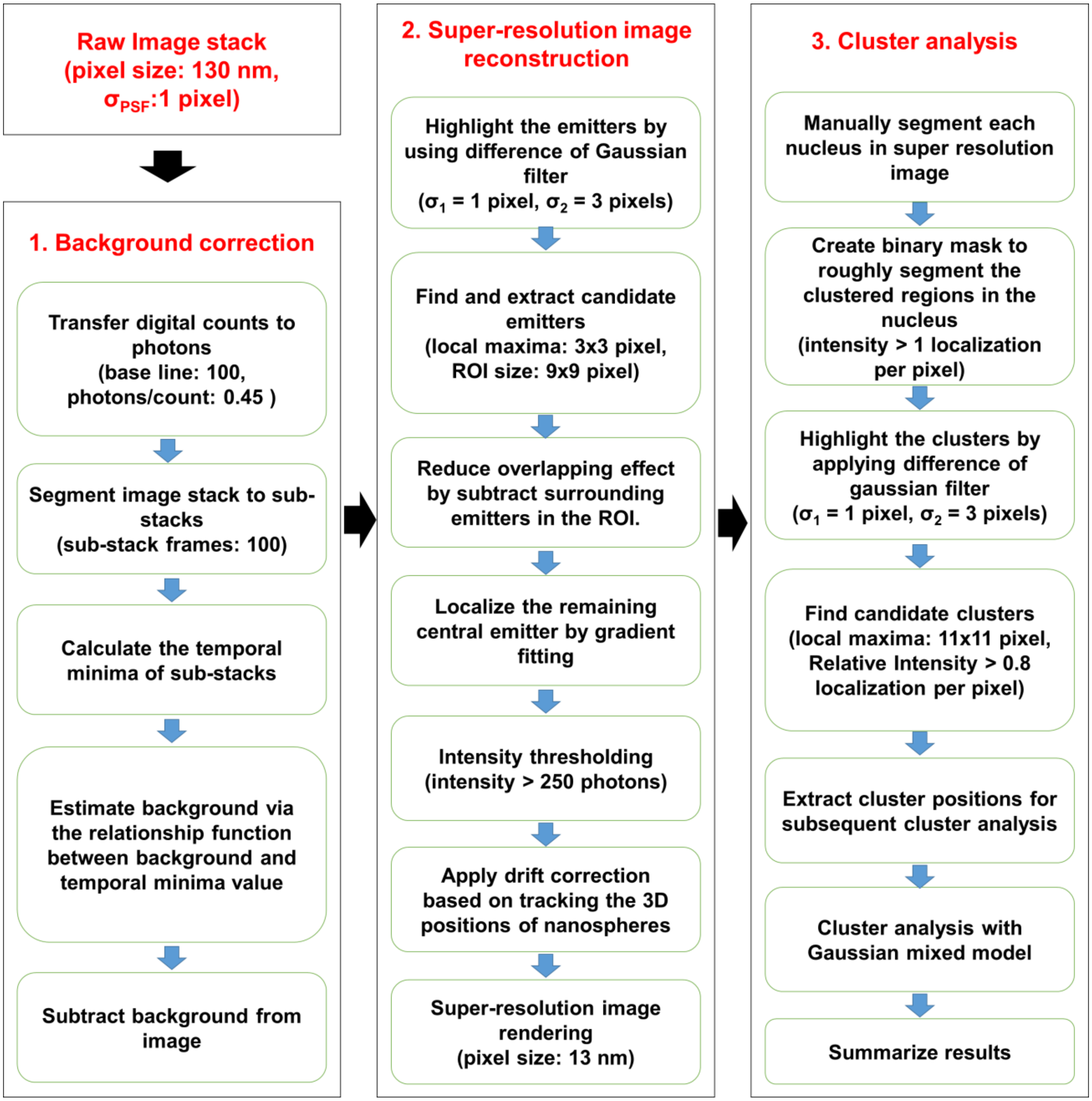


**Figure S10.** Flowchart to show the process and selected parameters for image reconstruction and cluster analysis.

**Supplementary Table**

**Table S1. Patient characteristics for human pathological tissue from surgical resection**

| Sample ID | Gender | Age | Group | Pathological stage of tissue imaged by PathSTORM | Most advanced pathological stage of the patient |
|---|---|---|---|---|---|
| 1 | female | 65 | Normal | normal | Diverticulosis |
| 2 | female | 51 | Normal | normal | Diverticulosis |
| 3 | female | 81 | Normal | normal | Diverticulosis |
| 4 | male | 80 | Normal | normal | Diverticulosis |
| 5 | male | 66 | Normal | normal | Diverticulosis |
| 6 | male | 65 | LGD/adenoma | LGD | LGD |
| 7 | female | 70 | LGD/adenoma | Tubular adenoma | Adenocarcinoma |
| 8 | male | 78 | LGD/adenoma | Adenoma | Sissle serrated adenoma |
| 9 | female | 76 | LGD/adenoma | Tubular adenoma | Tubular adenoma |
| 10 | male | 51 | LGD/adenoma | Tubular adenoma | Tubular adenoma |
| 11 | male | 59 | HGD | HGD | HGD |
| 12 | male | 81 | HGD | HGD | Tubular adenoma with focal HGD |
| 13* | female | 24 | HGD | Adenoma with HGD | Adenocarcinoma |
| 14 | female | 66 | HGD | HGD | Adenocarcinoma |
| 15 | female | 57 | Cancer | Adenocarcinoma | Adenocarcinoma |
| 16* | female | 24 | Cancer | Adenocarcinoma | Adenocarcinoma |
| 17 | female | 81 | Cancer | Adenocarcinoma | Adenocarcinoma |
| 18 | female | 80 | Cancer | Adenocarcinoma | Adenocarcinoma |
| 19 | female | 64 | Cancer | Adenocarcinoma | Adenocarcinoma |

* Sample #13 and #16 are from the same patient.
